# Membrane-induced 2D phase separation of focal adhesion proteins

**DOI:** 10.1101/2023.03.31.535113

**Authors:** Thomas Litschel, Charlotte F. Kelley, Xiaohang Cheng, Leon Babl, Naoko Mizuno, Lindsay B. Case, Petra Schwille

## Abstract

Focal adhesions form liquid-like assemblies around activated integrin receptors at the plasma membrane. Made up of hundreds of proteins, focal adhesions are dynamic structures which can assemble and disassemble quickly, withstand strong actomyosin-applied forces, and form highly stable complexes. How they achieve these flexible characteristics is not well understood. Here, we use recombinant focal adhesion proteins to reconstitute the core structural machinery *in vitro*, with the goal of understanding the underlying protein dynamics and interactions. We observe liquid-liquid phase separation of the core focal adhesion proteins talin and vinculin for a spectrum of conditions and in combination with several interaction partners. Intriguingly, we show that membrane binding triggers phase separation of these proteins on the membrane, which in turn induces the enrichment of integrin in the clusters. We also introduce a novel experimental setup to probe talin-membrane interactions down to the single protein level. Our results suggest that membrane composition triggers condensate assembly at the membrane, a regulatory mechanism which could widely apply to membrane-localized biomolecular condensates and provide a pathway of how spatial organization of lipids within the membrane can couple into the cytosol.

## Introduction

Phase separation driven by interactions between multivalent molecules can drive the formation of cellular compartments, termed biomolecular condensates. Condensates often exhibit liquid-like material properties, can contain hundreds of constituent molecules that are densely packed, and their formation is dependent on the phase separation of a few components. At the plasma membrane, protein-driven phase separation can promote receptor clustering and assembly of many adhesion and signaling complexes (*1, 2*). Due to the presence of the plasma membrane, membrane-associated condensates contain an intrinsic polarity.

Focal adhesions connect the actin cytoskeleton to the extracellular matrix, thereby regulating cell shape, migration, differentiation, and cellular responses to extracellular stimuli (*3*). Centered around activated integrin receptors in the plasma membrane, focal adhesions are highly dynamic assemblies made up of densely packed proteins (*4, 5*). These complexes can form and disassemble rapidly in response to cellular cues, or form stable, long-lasting connections between the extracellular matrix and cellular actomyosin networks. Focal adhesion proteins interact mainly through weak, often multivalent, interactions and can rapidly exchange between focal adhesions and the cytoplasm (*6-8*). Individual focal adhesions grow and mature over time and can fuse with each other (*9-11*). Consistent with these characteristics, recent studies demonstrate that focal adhesion adaptor proteins can undergo phase separation, suggesting that phase separation may contribute to integrin clustering, nascent focal adhesion assembly (*12, 13*), focal adhesion maturation (*14*), and focal adhesion disassembly (*15*). However, mature focal adhesions in cells have distinct vertical layers of protein localization above the membrane (*16*). How can phase separation of integrin receptors regulate protein organization 100 nm above the membrane surface, and *vice versa*? Identifying the underlying principles governing focal adhesion organization, composition, and dynamics is crucial for understanding the unique and varied roles these complexes play in cell adhesion.

Talin, a large (272 kDa) mechanosensitive scaffolding protein (Fig. 1A), is an ideal candidate for regulating both integrin clustering and protein organization above the membrane. Talin connects integrin receptors to the actin cytoskeleton, and is essential for focal adhesion formation (*17, 18*). Roughly 100 nm long, talin has a highly polarized orientation within focal adhesions; its N-terminus localizes with integrin receptors at the membrane while its C-terminus localizes to the site of stress fiber attachment. Talin spans the entirety of focal adhesion complexes and is the primary determinant of the nanoscale organization within focal adhesions (*19, 20*). Because of its key role in focal adhesion formation and the organization of focal adhesion nanostructure, we characterized the ability of talin to form condensates in vitro using purified recombinant proteins. We found that talin undergoes phase separation to form liquid-like condenates. Talin conformation is highly regulated and is maintained in an autoinhibited conformation in the cytoplasm (*20*). We therefore looked into interactions between talin and other core components of focal adhesions and find that talin activation regulates its condensation on the membrane. Interactions between talin and PI(4,5)P_2_ were sufficient to release authoinhibition and promote talin phase separation on membranes, and PI(4,5)P_2_ mediated clustering strengthens talin-membrane connections.

**Fig. 1.**
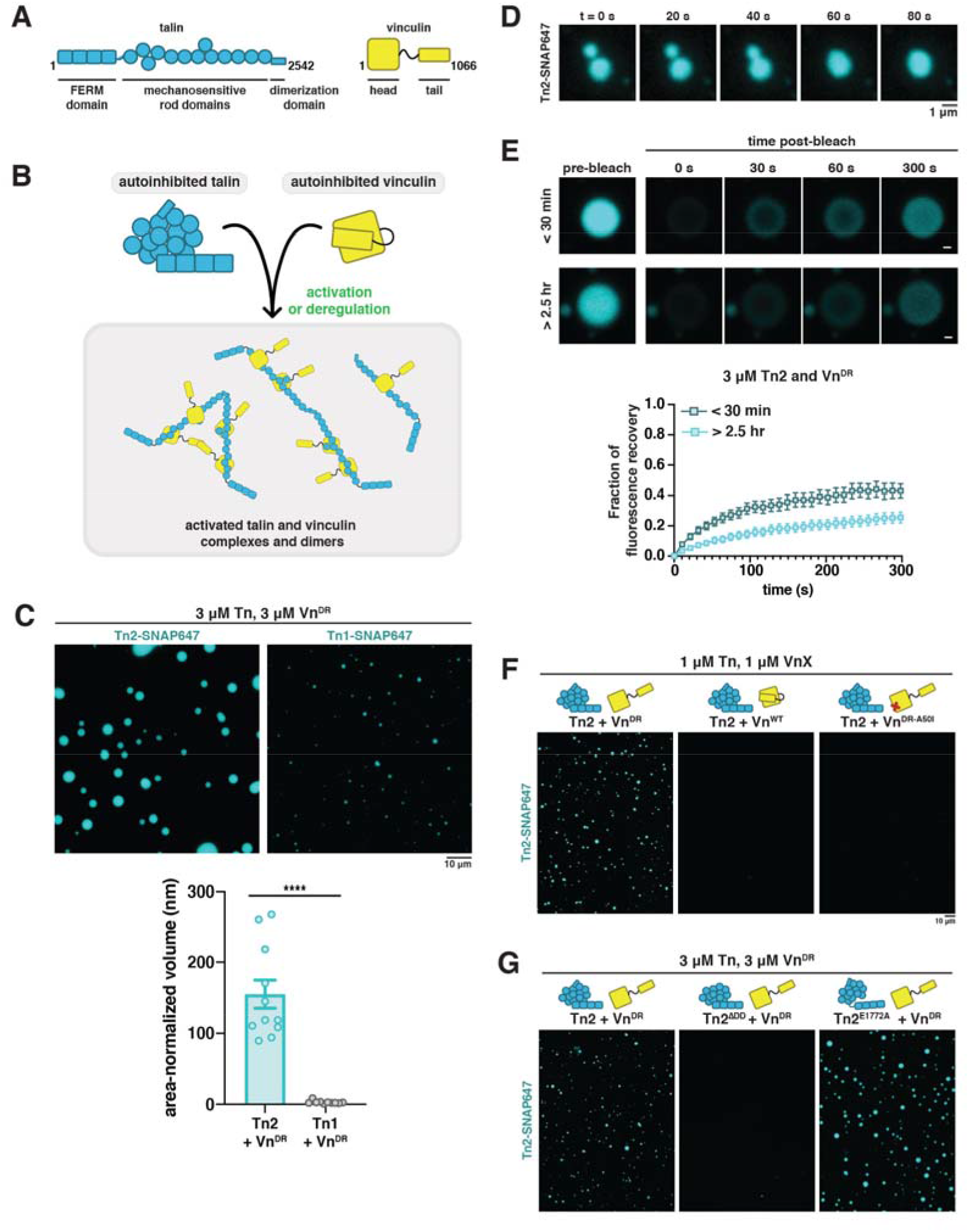
Talin undergoes liquid-liquid phase separation. (**A**) Domain schematics for talin and vinculin. **(B)** Protein autoinhibition needs to be released or removed in order for complexes to form. **(C)** Talin2 (Tn2) forms droplets when incubated with deregulated vinculin (Vn^DR^) under crowded conditions at room temperature. Talin1 (Tn1) forms smaller droplets under the same conditions. **(D)** Fusion of Tn2 and Vn^DR^ droplets demonstrate liquid-like behavior. **(E)** Photobleaching of Tn2-Vn^DR^ droplets indicate fluorescence recovery. Recovery slows as the droplets age. **(F)** Tn2-Vn^DR^ droplet formation can be disrupted by mutation of the talin binding site in vinculin (A50I). **(G)** Removing the talin dimerization domain (Tn2^ΔDD^) reduces droplet formation, while reducing talin autoinhibitory interactions between the rod and head regions (E1772A mutation) increases phase separation. All conditions for (C,F,G) were repeated in triplicate, and imaged after incubation at room temperature for one hour. See supplementary Figure S6 for quantification. Purified proteins were mixed in the following buffer (10 mM imidazole, 50 mM KCl, 1 mM MgCl_2_, 1 mM EGTA, 0.2 mM ATP, pH 7.5) supplemented with 15 mM glucose, 20 μg/mL catalase, 100 μg/mL glucose oxidase, 1 mM DTT and 0.25% methyl-cellulose (4000 cp). Error bars represent SEM. **** p > .0001

## Results

We first looked into interactions between recombinant full-length talin and recombinant vinculin (Fig. 1). Vinculin (Fig. 1A) interacts directly with talin and actin, and is recruited to growing focal adhesions to regulate mechanostransduction. Both talin and vinculin need to be relieved of their autoinhibited conformation in order to interact (Fig 1B). To focus on the activation of talin, we designed a deregulated vinculin (Vn^DR^) by disrupting the two autoinhibitory interactions between the vinculin head and vinculin tail domains (*21, 22*). Removing these inhibitory interaction results in the formation of Tn-Vn droplets (Fig. 1C). Vertabrates contain two talin genes, Tn1 and Tn2, which are 76% identical but not functionally redundant (*23*). We compared the two talin isoforms, Tn1 and Tn2, mixed with Vn^DR^ and used methyl cellulose as a crowder to imitate conditions in the cytoplasm. Under these conditions, Tn2 formed micrometer sized droplets, while Tn1 formed smaller droplets and produced a drastically lower volume of phase separated material (Fig. 1C), even at higher protein concentrations (Fig. S2A). It is possible that Tn1 needs further activation, or an additional binding partner, to reach the level of phase separation observed with Tn2-Vn^DR^. DeePhase, a predictor of protein phase behavior based on amino acid sequence, did not find talin1 any less likely to phase separate than talin2 (*24*). Therefore, the apparently different thresholds for phase separation of talin1 and talin2 is likely due to small differences in the strength of autoinhibitory, intramolecular interactions.

We confirmed the liquid-like nature of the talin-vinculin droplets by observing the coalescence of multiple droplets (Movie1) and through fluorescence recovery after photobleaching (FRAP) measurements (Fig. 1D,E and Fig. S2B,C). Recovery speed decreased over time for Tn2 droplets, suggesting talin condensates mature and gradually become more viscous over time (*25*). The volume of protein-rich phase depends on both protein and crowder concentration, and is salt-sensitive (Fig. S4, Fig. S5). Tn2 did not phase separate with wild-type Vn or with Vn^DR-A50I^, a deregulated vinculin mutant unable to bind to talin (Fig. 1F, Fig. S6A), indicating that specific talin-vinculin interactions are required for phase separation. The talin dimerization domain is also required for droplet formation, and a point mutation reducing interactions between the talin head and rod domains (Tn2^E1772A^) increased the volume of the protein-rich phase, suggesting talin autoinhibition lowers its propensity to phase separate (Fig. 1G, Fig. S6B). However, neither Tn2^E1772A^ alone nor Tn with Vn^D1^, the vinculin fragment containing the talin binding site, were sufficient for phase separation (Fig. S6B).

The Beta integrin cytoplasmic domain binds directly to the talin FERM domain (*26*) and integrin binding promotes talin activation. To test whether talin activation by integrin binding would promote phase separation, we synthesized a fluorescently-labeled peptide of the β1D integrin cytoplasmic tail sequence (Fig. 2A). The Tn2-integrin β1D interaction is the strongest of the characterized talin-integrin receptor interactions (∼30 μM) (*27*). Tn2 formed liquid-like droplets with the β1D peptide alone under crowded conditions (Fig. 2B), without an additional activating binding partner such as Vn^DR^, suggesting that β1D is sufficient to activate Tn2. Phase separation was dependent on the concentration of the β1D peptide, which was required in excess (Fig. S7). No phase separation was observed when Tn1 and β1D were mixed, and mutation of the residue responsible for the increased affinity of talin2 for β1D domain (Tn2^S339L^) severely reduced the volume of the protein-rich phase (Fig. 2B, Fig. S8) (*28*). β1D is concentrated within Tn2 condensates, suggesting that talin phase separation could be sufficient to promote integrin clustering (*29*). Interestingly, in these droplets Tn2 recovery after photobleaching was reduced compared to Tn2-Vn^DR^ droplets, while β1D rapidly recovered (Fig. 2C), suggesting that dynamics within Tn2-based condensates can vary based on binding partners. Wild-type Vn was recruited to Tn2-β1D droplets, but reduced the total volume of the protein-rich phase, indicating that vinculin may limit the extent of talin-induced integrin clustering (Fig. 2D, Fig. S9).

**Fig. 2.**
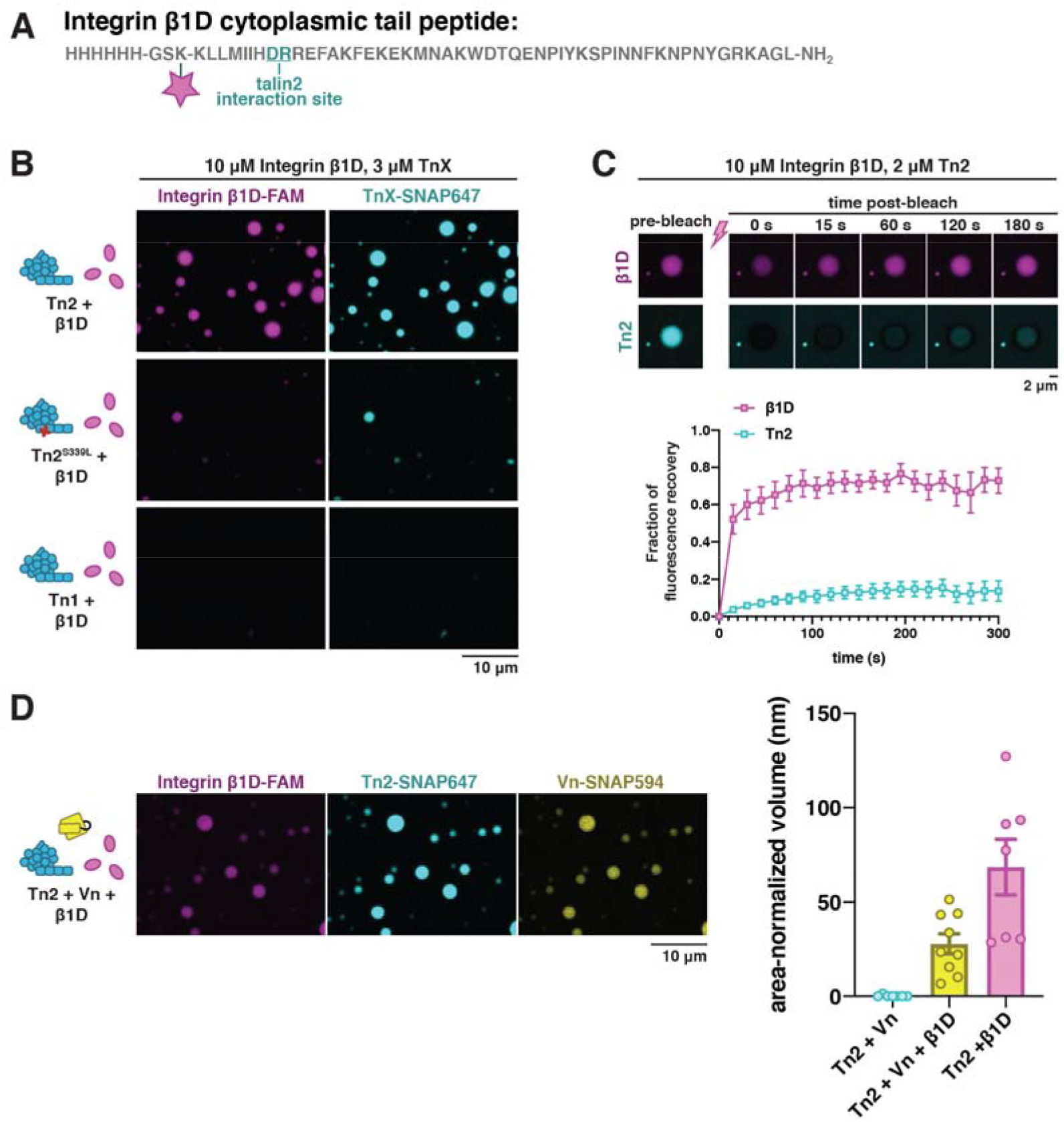
Integrin β1D drives phase separation of talin. **(A)** Synthetic Integrin β1D cytoplasmic tail peptide labeled with carboxyfluorescein(FAM). **(B)** The β1D peptide specifically drives phase separation of Tn2, not Tn1, under crowded conditions at room temperature. Phase separation can be disrupted by a mutation in the β1D binding site. **(C)** Photobleaching of β1D-Tn2 droplets indicate that Tn2 and the β1D peptide reveal different recovery rates. **(D)** Wild-type vinculin is also recruited to Tn2-β1D droplets, but slightly decreases the amount of phase separated material. All conditions for (B,D) were repeated in triplicate, and imaged after incubation at room temperature for one hour. Purified proteins were mixed in the following buffer (10 mM imidazole, 50 mM KCl, 1 mM MgCl_2_, 1 mM EGTA, 0.2 mM ATP, pH 7.5) supplemented with 15 mM glucose, 20 μg/mL catalase, 100 μg/mL glucose oxidase, 1 mM DTT and 0.25% methyl-cellulose (4000 cp). Error bars represent SEM.

Together, these results sugest that activation of Tn by either Vn^DR^ or β1D is sufficient to promote talin phase separation. We further found that inducing talin activation with salt also promotes talin phase separation (Fig. S10). As previously shown, salt can artificially induce talin activation in vitro by disrupting weak interactions between rod domains (*20*). Thus, we conclude that releasing talin autoinhibition consistently leads to its phase separation.

While both activated vinculin and β1 integrin are predicted to release talin autoinhibition, they are unlikely to be the primary talin activator during focal adhesion initiation and assembly. Instead, the phosphoinositide PI(4,5)P_2_ plays a major role in regulating talin localization and activation at the plasma membrane, and PI(4,5)P_2_ is necessary for proper formation of functional focal adhesions (*30*). Binding to PI(4,5)P_2_ triggers a shift from a globular, inactive conformation to an open, extended conformation, in which talin is able to recruit vinculin and actin to membrane surfaces (*20, 22, 31*). Therefore, we wished to test whether PI(4,5)P_2_-containing membranes can also trigger phase separation by releasing talin autoinhibition. We extended our microscopy assay with phase separating protein solutions to include lipids as additional binding partners in the reaction mix. To this end, we prepared relatively homogeneous dispersions of small unilamellar vesicles (SUVs) (<100 nm in diameter) and mixed them with Tn2 under physiological conditions with a crowding agent. Excitingly, we observed phase separation of Tn2 when combined with a dispersion of SUVs with PI(4,5)P_2_-rich membranes, but not for the control with SUVs that do not contain PI(4,5)P_2_ (Fig. 3A, Fig. S11). Phase separation of Tn2 was enhanced by including either the β1D peptide or Vn^DR^. Tn1 did not form droplets when mixed with 20% PI(4,5)P_2_ SUVs (Fig. 3B), consistent with the lower affinity of Tn1 for PI(4,5)P_2_ (*32*). Interestingly, droplets with Tn2, β1D, and SUVs were much more viscous, clumping together instead of forming spherical droplets (Fig. S12). Even more strikingly, these condensates did not require crowded conditions to form, indicating a stronger tendency to phase separate than with either talin activator independently.

**Fig. 3.**
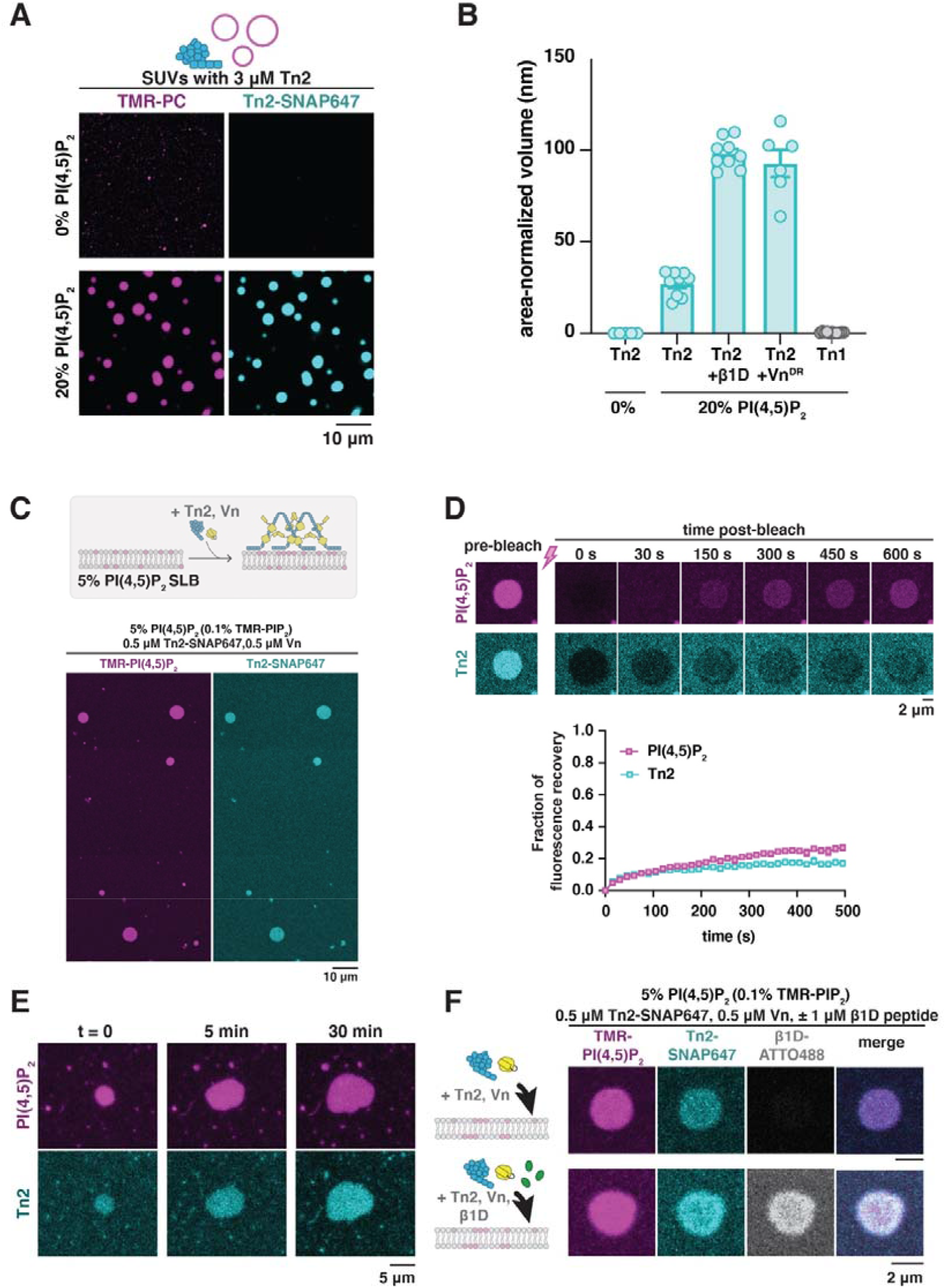
Membrane-bound talin phase separates. (**A**) Tn2 forms droplets in the presence of PI(4,5)P_2_-rich vesicles under crowded conditions, but not in the absence of PI(4,5)P_2_. Top: SUVs consisting of 75% POPC, 15% POPE, 10% POPS, Bottom: SUVs consisting of 55% POPC, 20% PI(4,5)P_2_, 15% POPE, 10% POPS. Note that vesicle sizes are below the resolution limit; visible structures are liquid condensates. **(B)** Phase separation in the presence of vesicles is enhanced by the addition of phase separation promoting talin binding partners, β1D and Vn^DR^. **(C)** Talin forms 2D clusters on PI(4,5)P_2_-rich supported lipid bilayers (SLBs). **(D)** Tn2-PI(4,5)P_2_ clusters recover fluorescence slowly after photobleaching. **(E)** Tn2-PI(4,5)P_2_ clusters change shape and grow over time. **(F)** Integrin β1D peptide is also recruited to Tn2-PI(4,5)P_2_ clusters on the membrane. All lipid-based experiments were carried out in the following buffer (10 mM HEPES pH 7.5, 100 mM NaCl), either with 0.25% methyl cellulose (vesicle experiments) or without crowding reagent (SLB experiments).

Next, we added Tn2 to supported lipid bilayers (SLBs) containing 5% PI(4,5)P_2_, to investigate the phase separation of talin on membranes more closely resembling those of living cells. When added together with wild-type Vn, micron and sub-micron two-dimensional (2D) circular Tn clusters form immediately (<5 min), colocalizing with labeled TMR-PI(4,5)P_2_ clusters in the membrane (Fig. 3C). These clusters require a minimum amount of PI(4,5)P_2_ in the membrane to form (Fig. S13), and recover partially after photobleaching (Fig. 3D). In addition, clustering requires the talin dimerization domain, suggesting a mechanism analogous to 3D droplet formation in solution under crowded conditions (Fig. S14). The clusters can be seen to grow over time, with fluctuating boundaries (Fig. 3E). These observations suggest that while the membrane clusters may not be as fluid as their soluble counterparts, neither are they static structures. Interestingly, in the presence of polymerizing actin, the clusters take on more irregular shapes, but do not seem to recruit actin filaments or mediate polymerization in any way (Fig. S15). The β1D peptide is strongly recruited to the clusters, at a much lower talin-to-peptide ratio than required in solution under crowded conditions (Fig. 3F). While clustering of transmembrane, alpha-beta subunits of integrin receptors is undoubtedly a more complex process, the ability of membrane-bound talin to recruit the β1D peptide to the membrane and form 2D, phase separated micro-domains indicates that it would have a similar effect on full-length integrin receptors. Integrin-talin clusters are thought to be precursors of FAs, and are dependent on cellular PI(4,5)P_2_ levels, supporting the idea that PI(4,5)P_2_-talin interactions play a role in integrin clustering and FA assembly (*29*).

To better understand the interactions between talin and the membrane, we developed an assay to measure the strength of single Tn2-PI(4,5)P_2_ interactions using optical tweezers (Fig. 4A). We coated one type of micron-sized beads with Tn2 via a DNA handle and biotin-neutravidin interaction and a second type of beads with PI(4,5)P_2_. A dual trap optical tweezer setup was used to catch one of each bead and manipulate their positions to measure interactions between individual proteins and the membrane.

**Fig. 4.**
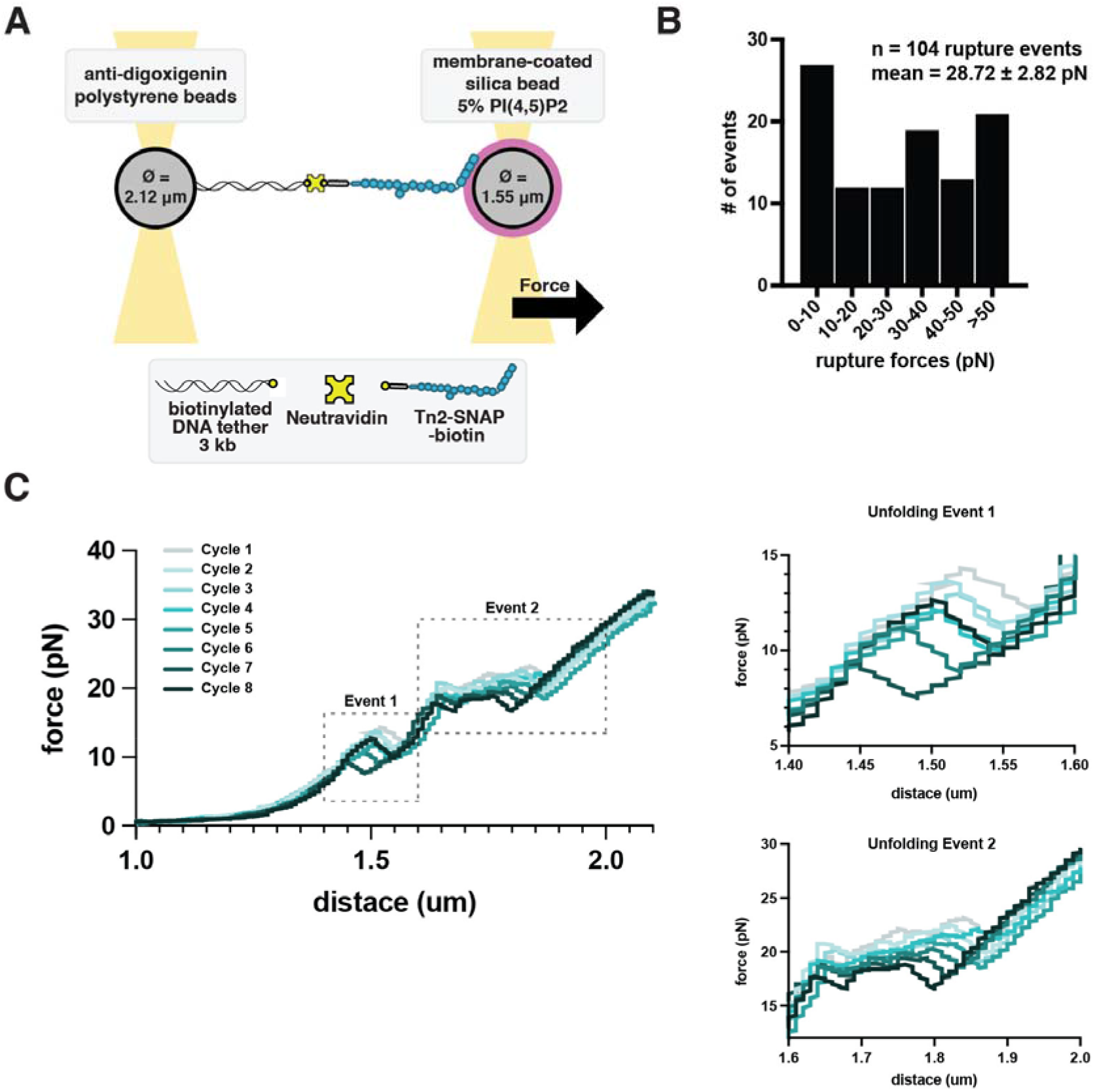
Clustering strengthens talin-membrane connections. **(A)** Optical trap set up for measuring single talin-membrane interactions. **(B)** Rupture force distribution for interaction of Tn2^ΔDD^ with PI(4,5)P_2_ bilayer. We measured the forces at which the protein detaches from the membrane. **(C)** Tn2^ΔDD^ force curve pre-rupture showing protein unfolding. We can identify two unfolding events, which matches previous reports for the isolated talin rod domain. All optical-trap experiments were carried out in the following buffer (10 mM HEPES pH 7.5, 100 mM NaCl).

We found Tn2 most often formed multiple tethers between the beads and interactions were strong enough to frequently pull the membrane-coated bead out of its optical trap. Occasionally we were able to record multiple, sequential rupture events (Fig. S17). We hypothesized that the strength of talin-membrane interactions may be due to talin’s propensity to phase separate when membrane bound. To test this, we instead used Tn2^ΔDD^, thereby retaining the membrane affinity and mechanosensitivity of full-length Tn2, while eliminating potential clustering on the membrane. Indeed, measurements for Tn2^ΔDD^ samples resembled models of force curves of single proteins, and thus also allowed more thorough quantification. Rupture forces for Tn2^ΔDD^-membrane interactions ranged from 1 to 60 pN, with a mean rupture force of 28.72 (n = 104 rupture events) (Fig. 4B). Excitingly, these results clearly show that Tn2-membrane interactions can withstand forces within the range at which talin rod domains unfold. In fact, we were able to observe consecutive unfolding/refolding cycles, with two consistent and reversible unfolding events (Fig. 4C). These results confirm that the talin rod domains display similar mechanosensitive behavior in the context of the full-length, membrane-bound protein.

## Discussion

Strong evidence points to talin acting as a master regulator of focal adhesion assembly and organization. We have shown in this study that activation of talin through multiple pathways triggers the formation of LLPS condensates. Focal adhesions show liquid-like properties in cells, and share many characteristics with confirmed biomolecular condensates. However, focal adhesions naturally are highly anisotropic: they only exhibit these liquid-like properties in two dimensions laterally along the membrane, but show a highly structured organization orthogonal to the membrane (*16*) in large parts due to talin’s role as a core structural scaffold (*19*). Our in vitro experiments suggest that this anisotropicity could be explained by lipid-regulated phase separation, thus restricting a typically 3-dimensional behavior to two dimensions.

Other recent work has shown the wetting of protein condensates on membranes (*33-35*), and a lowering of the phase separation threshold by membrane binding (*36-38*) or clustering of transmembrane proteins (*39-41*). However, here we propose a new regulatory mechanism in which lipids act as an interaction partner required for the condensation of proteins, which suggests a truly 2-dimensional protein phase separation. Hence this gives way for a more structured organization of focal adhesions in the remaining third dimension, i.e. into the cytosol.

Biomolecular condensates have been identified in a range of physical states; varying greatly in terms of dynamics, stability, life-span, and function. Talin condensates in solution become more viscous over time, and protein turnover varies depending on components and composition. Interestingly, the recovery of Tn2 and PI(4,5)P_2_ within 2D membrane-associated clusters is fairly slow with a high immobile fraction, suggesting these condensates are more similar to bioreactive gels than to other highly fluid membrane-bound condensates. This would be consistent with talin’s inability to diffuse laterally along the membrane surface in cells. The mechanosensitive nature of focal adhesions may require this increased viscosity, in order to withstand forces applied through actomyosin contraction. A similar explanation has been offered for the gel-like state of centrosomal condensates, which must withstand forces applied by the spindle during mitosis (*42*).

We further demonstrate that individual talin-membrane interactions are strong enough to withstand forces capable of inducing rod domain unfolding events. Our measurements of the wild-type Tn2 also suggest that the ability to dimerize strengthens talin-membrane interactions beyond the sensitivity of the current experimental system, which is specifically designed to measure binary interactions. Further application of these methods could be used to reach a deeper understanding of the role of both talin-talin and talin-membrane interactions in focal adhesion mechanosensation. Our results suggest that interactions between talin and PI(4,5)P_2_ are likely necessary for talin to resist high mechanical forces within focal adhesions (up to 10 pN per molecule (*43*)), since the talin-integrin linkage is a low affinity interaction (30 – 500 μM) (*27*).

As more biomolecular condensates are identified in diverse cellular processes, it is becoming clear that interactions with membranes play an important role in their formation, function and regulation (*44-49*). Until now, lipid bilayers have been viewed mostly as passive surfaces which can localize or increase phase separate through physical effects. Here, we argue that specific lipids can play catalytic roles in regulating protein phase separation; PI(4,5)P_2_ not only concentrates talin at the membrane, but also releases talin autoinhibition. By activating talin, PI(4,5)P_2_ likely triggers phase separation and enables the recruitment of other adhesion proteins, thereby initiating a crucial step in focal adhesion assembly. PI(4,5)P_2_ binds and regulates additional focal adhesion adaptor proteins, including FAK and vinculin, suggesting lipids could coordinate the recruitment and phase separation of additional components. Our experiments suggest that spatial organization on the level of lipids could have structural effects that reach far into the cytoplasm.

## Acknowledgements

We thank Michaela Schaper for help by cloning plasmids and we thank Stefan Übel for synthesizing the β1D integrin peptide. This work is part of the MaxSynBio consortium, which is jointly funded by the Federal Ministry of Education and Research of Germany and the Max Planck Society. C.F.K. is a recipient of the Humboldt Research Fellowship for Postdoctoral Researchers and has received funding from the European Union’s Horizon 2020 research and innovation program under the Marie Sklodowska-Curie grant agreement No 794162. N.M. acknowledges the Boehringer Ingelheim Foundation Plus 3 Program, and the European Research Council (ERC-CoG, 724209).

