## Supplementary Figures for "Membrane-induced 2D phase separation of focal adhesion proteins"

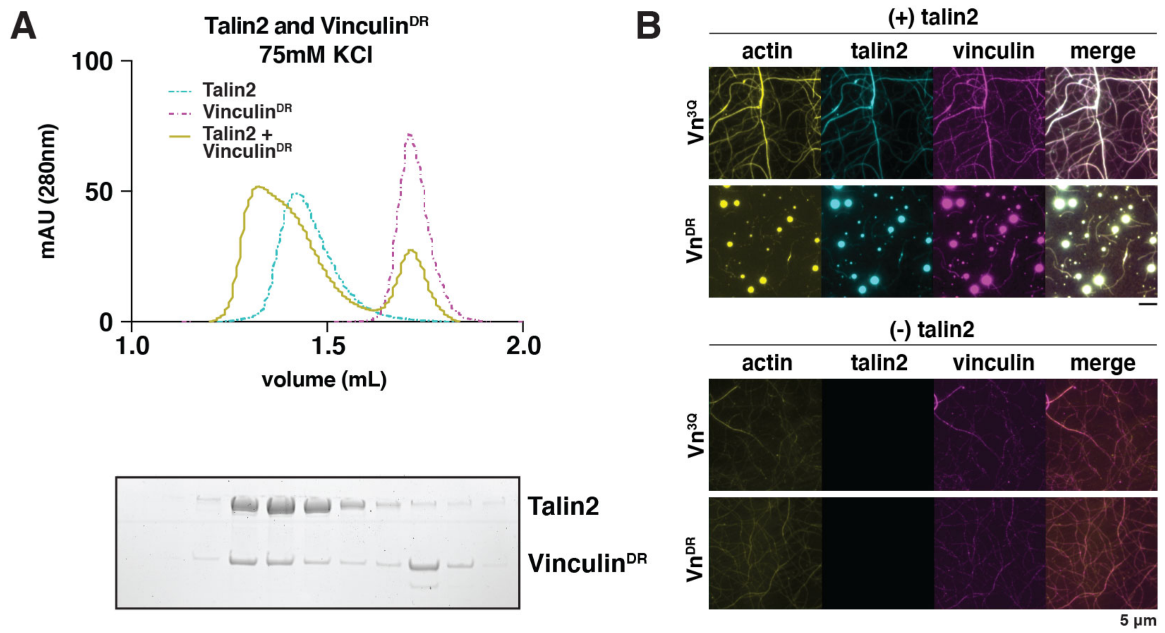


**S1. Vn$^{DR}$ can bind to Tn2 in low salt conditions, and requires Tn2 for phase separation. (A)** Tn2 and Vn$^{DR}$ (both 5 µM) reconstitution assay using size-exclusion chromatography (SEC). Chromatograms and SDS-PAGE indicate the elution profiles of the proteins alone and in combination for (top) and Tn2,Vn$^{DR}$ (bottom) in 20 mM HEPES pH 7.8, 75 mM KCl, 1 mM EDTA, 3 mM β-mercaptoethanol. **(B)** The Vn$^{3Q}$ mutation disrupts one of two vinculin head-tail autoinhibitory interactions, while Vn$^{DR}$ has mutations at both autoinhibitory interfaces. Representative three-color TIRF microscopy images of 1.5 µM SNAP-tag-labeled FA proteins and 1 µM actin (10% actin-ATTO488) added to TIRFm buffer (10 mM imidazole, 50 mM KCl, 1 mM $MgCl_2$, 1 mM EGTA, 0.2 mM ATP, pH 7.5) supplemented with 15 mM glucose, 20 µg/mL catalase, 100 µg/mL glucose oxidase, 1 mM DTT and 0.25% methyl-cellulose (4000 cp). Images acquired after 30 min of actin polymerization. Vn$^{3Q}$ and Vn$^{DR}$ do not show obvious differences when incubated with actin alone, but in the presence of Tn, Vn$^{DR}$ results in droplets while Vn$^{3Q}$ results in actin bundling. This suggests that both autoinhibitory interactions between the vinculin head and tail domains must be disrupted to trigger phase separate with talin. Scale bar is 5 µm.

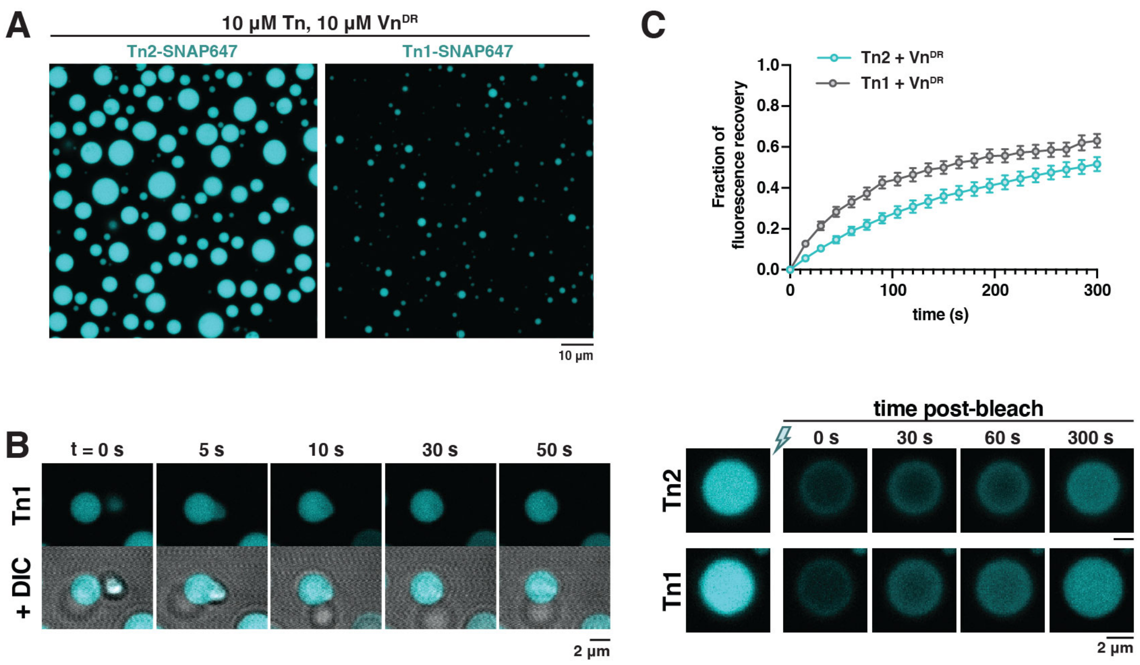


**S2. Talin1 requires higher concentrations to phase separate with Vn$^{DR}$. (A)** Representative image of Tn2-SNAP and Tn1-SNAP (10 µM) droplets that form upon mixing with Vn$^{DR}$ (10 µM) in the presence of a crowding agent. Each sample was done in triplicate, and incubated for one hour before imaging. All samples were mixed in the following buffer (10 mM imidazole, 50 mM KCl, 1 mM $MgCl_2$, 1 mM EGTA, 0.2 mM ATP, pH 7.5) supplemented with 15 mM glucose, 20 µg/mL catalase, 100 µg/mL glucose oxidase, 1 mM DTT and 0.25% methyl-cellulose (4000 cp). **(B)** Tn1-Vn$^{DR}$ droplets have liquid-like properties, and return to a spherical shape after fusing. **(C)** Fluorescence recovery after photobleaching of Tn1-Vn$^{DR}$ and Tn2-Vn$^{DR}$ droplets. Bleaching was carried out 15-30 minutes after initial droplet formation. Error bars represent SEM; n = 12 droplets for Tn1,Vn$^{DR}$, n= 6 droplets for Tn2,Vn$^{DR}$. Scale bars are indicated for each figure panel.

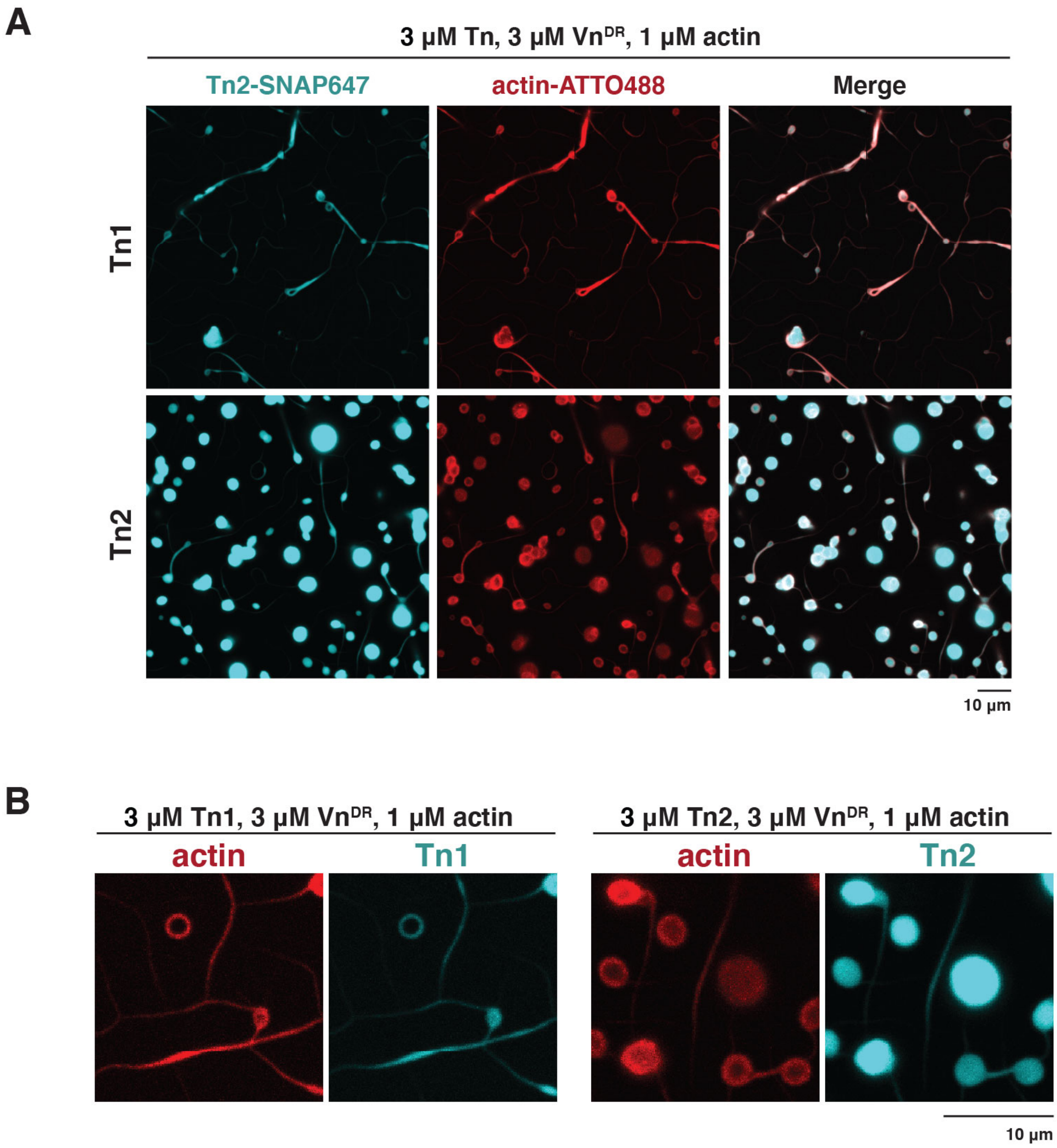


**S3. Talin2 forms more droplets in the presence of actin than talin1.** Representative three-color TIRF microscopy images of 3 μM SNAP-tag-labeled FA proteins and 1 μM G-actin (10% actin-ATTO488) in the following buffer (10 mM imidazole, 50 mM KCl, 1 mM $MgCl_2$, 1 mM EGTA, 0.2 mM ATP, pH 7.5) supplemented with 15 mM glucose, 20 μg/mL catalase, 100 μg/mL glucose oxidase, 1 mM DTT and 0.25% methyl-cellulose (4000 cp). Images acquired after 30 min of polymerization. **(A)** Tn1 with $Vn^{DR}$ bundles actin filaments, occasionally forming droplet-like spots. Tn2 with $Vn^{DR}$ forms actin-containing droplets, sometimes connected to actin bundles. **(B)** Zoomed-in view of Tn-$Vn^{DR}$-actin structures. Experiment done in triplicate. Scale bars are 10 μm.

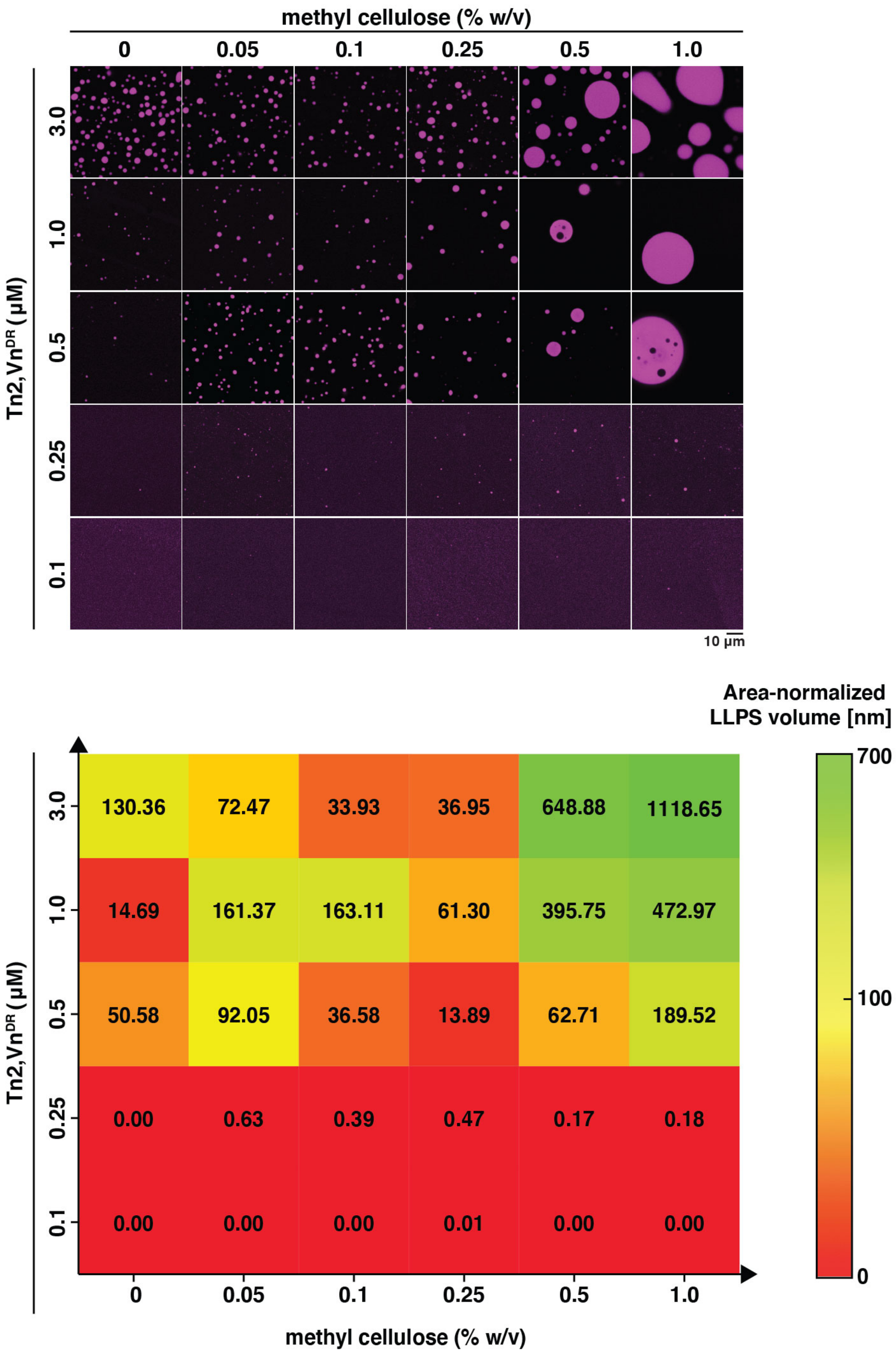


**S4. Phase separation of talin is dependent on protein and crowding agent concentration.** Representative images of droplet formation for varying concentrations of Tn2 and $Vn^{DR}$, with varying amounts of crowding agent (methyl cellulose – 4000 cp). These experiments were carried out in a 196-well plate, with slightly different results from those observed in channel-slides. Still, the dependence on concentration is consistent with all other experiments, though the threshold for droplet formation varies between imaging set ups. Heat map represents quantification of area-normalized LLPS volume for three independent samples for each condition. Scale bar is 10 $\mu$m.

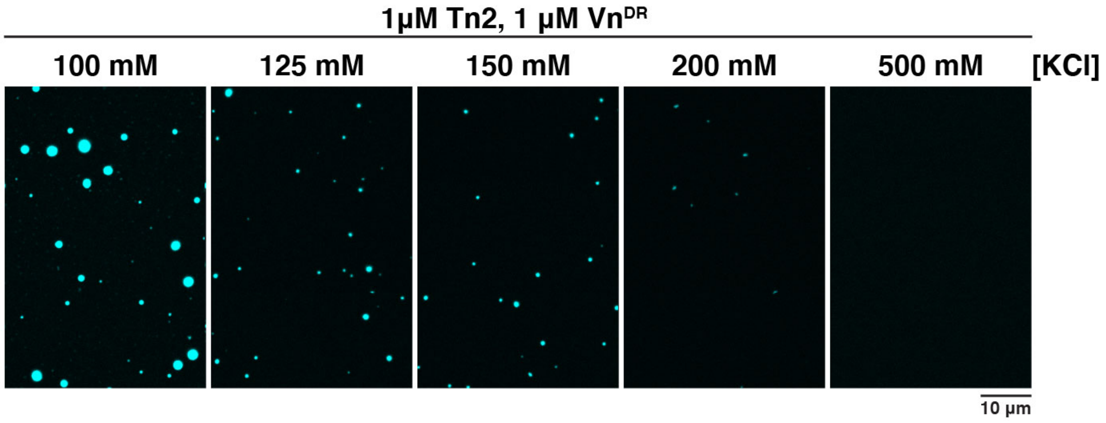


**S5. Phase separation of talin is salt sensitive.** The amount of phase separated material decreases with increasing amount of salt. Above 150 mM KCl, very few droplets are observed. Experiment done in triplicate, each sample was incubated for one hour in the following buffer (10 mM imidazole, X mM KCl, 1 mM $MgCl_2$, 1 mM EGTA, 0.2 mM ATP, pH 7.5) before imaging. Scale bar is 10 μm.

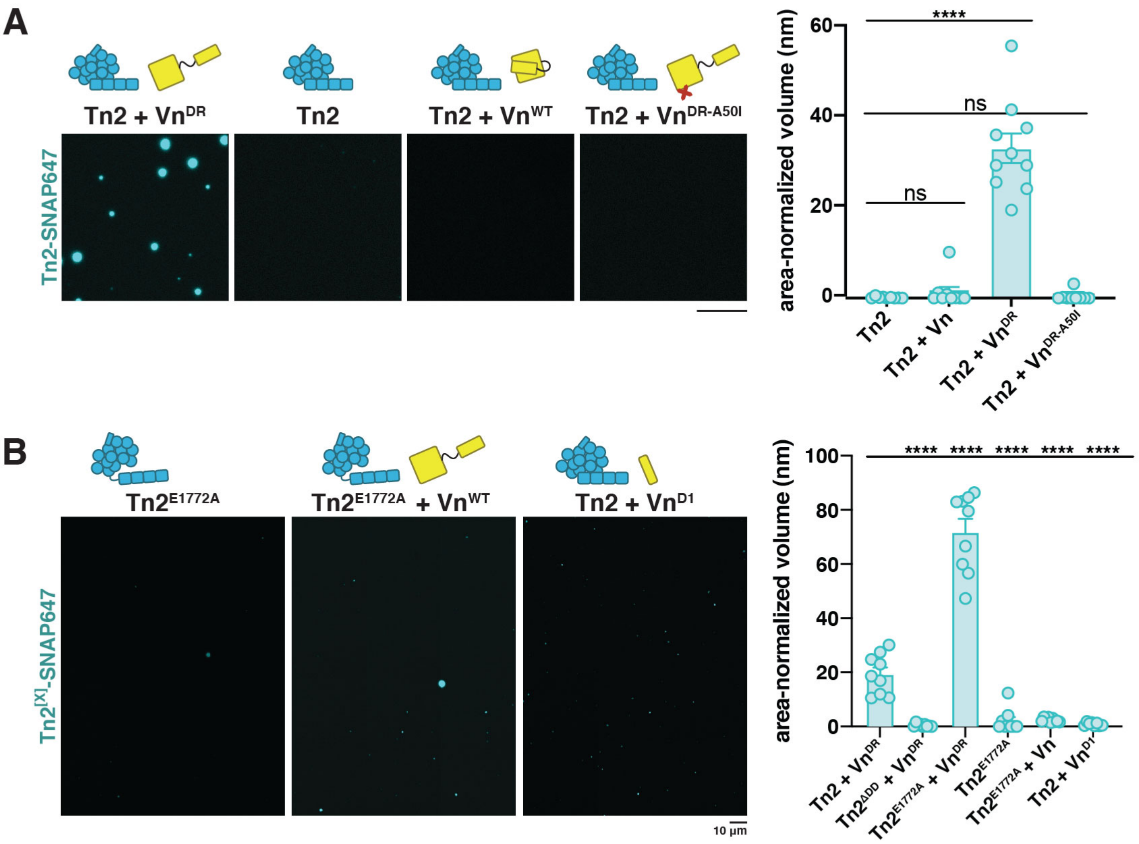


**S6. Phase separation of talin requires full-length, deregulated vinculin and the talin dimerization domain.** Additional conditions for Fig. 1F (A) and 1G (B), and quantification of the volume of phase separated material from confocal z-stacks. Each condition was performed in triplicate, multiple images from each sample were then pooled for quantification. Error bars represent SEM, scale bar is 10 $\mu$m.

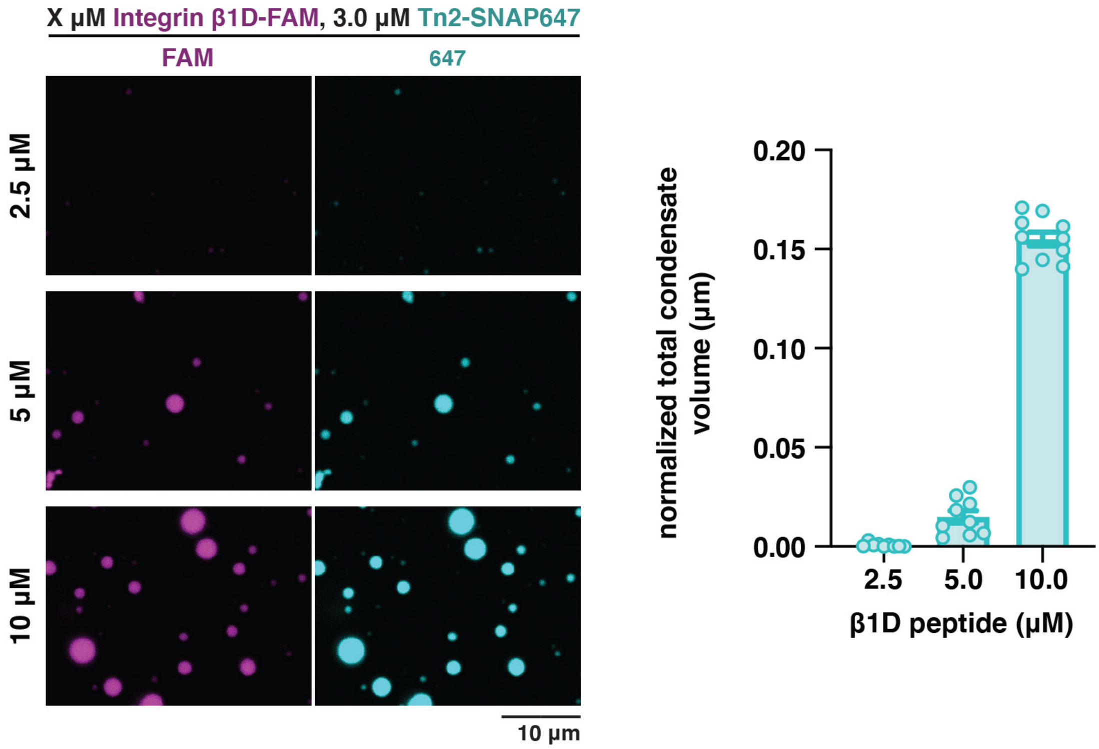


**S7. Phase separation of talin with β1D peptide is concentration dependent.** Representative confocal microscopy images of Tn2-β1D droplets formed at increasing concentrations of β1D integrin peptide. The amount of total condensate volume is quantified for the varying concentrations from confocal z-stacks. Tn2 was mixed with β1D integrin peptide and incubated for 1 hour before imaging. Each condition was performed in triplicate, multiple images from each sample were then pooled for quantification. Error bars represent SEM, scale bar is 10 μm.

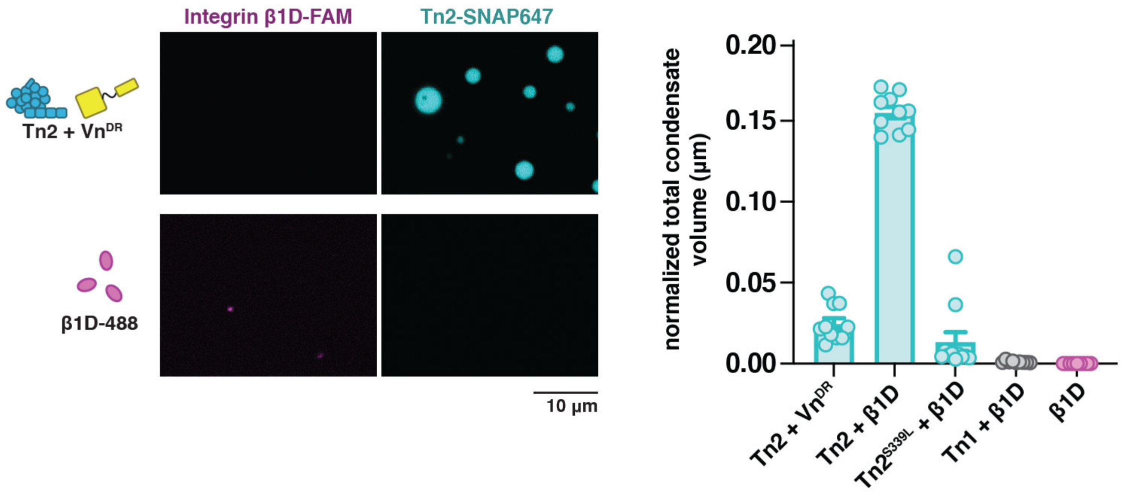


**S8. Integrin β1D drives phase separation of Tn2, not Tn1.** Additional conditions and quantification from Fig. 2B. Tn2-β1D droplet formation requires the residue responsible for Tn2's higher affinity for the β1D integrin receptor, indicating a specific effect. Additionally, Tn1 does not form droplets with the β1D peptide, consistent with the importance of the talin2 S339L residue. 3 μM TnX was mixed with 10 μM β1D integrin peptide and incubated for 1 hour before imaging. Each condition was performed in triplicate, multiple images from each sample were then pooled for the quantification. Error bars represent SEM, scale bar is 10 μm.

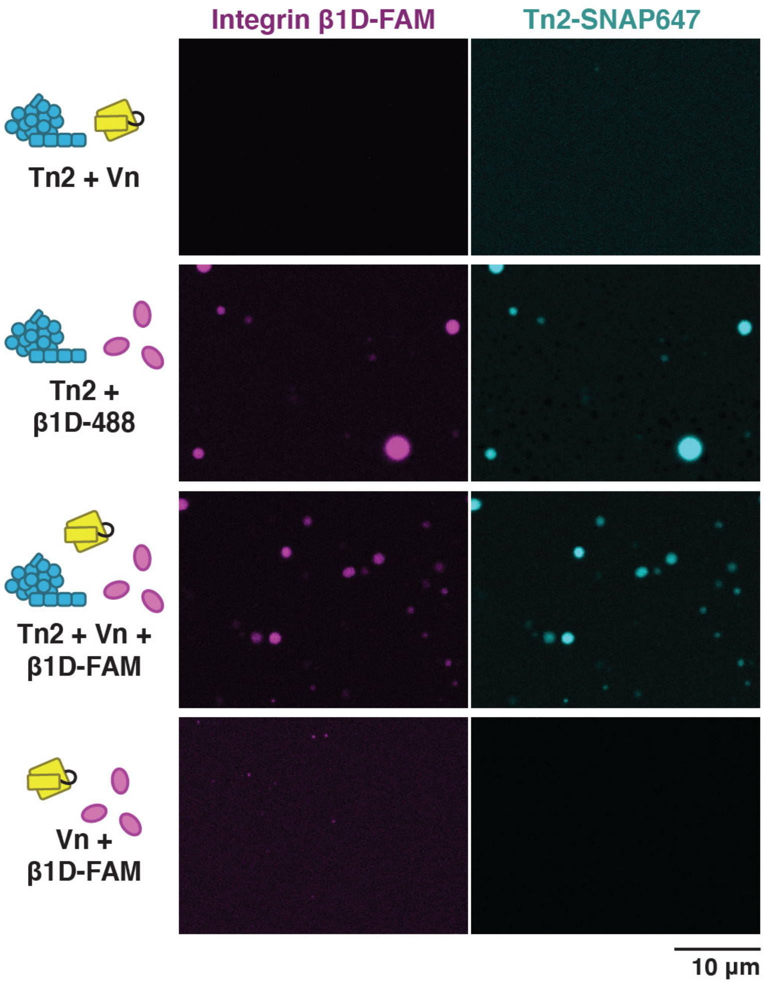


**S9. Vinculin reduces phase separation of Tn2-β1D, and does not phase separate with β1D alone.** Representative images for data displayed in Fig. 2D. 3 μM Tn2 and/or 3 μM Vn were mixed with 10 μM β1D integrin peptide and incubated for 1 hour before imaging. Scale bar is indicated in the figure. Scale bar is 10 μm.

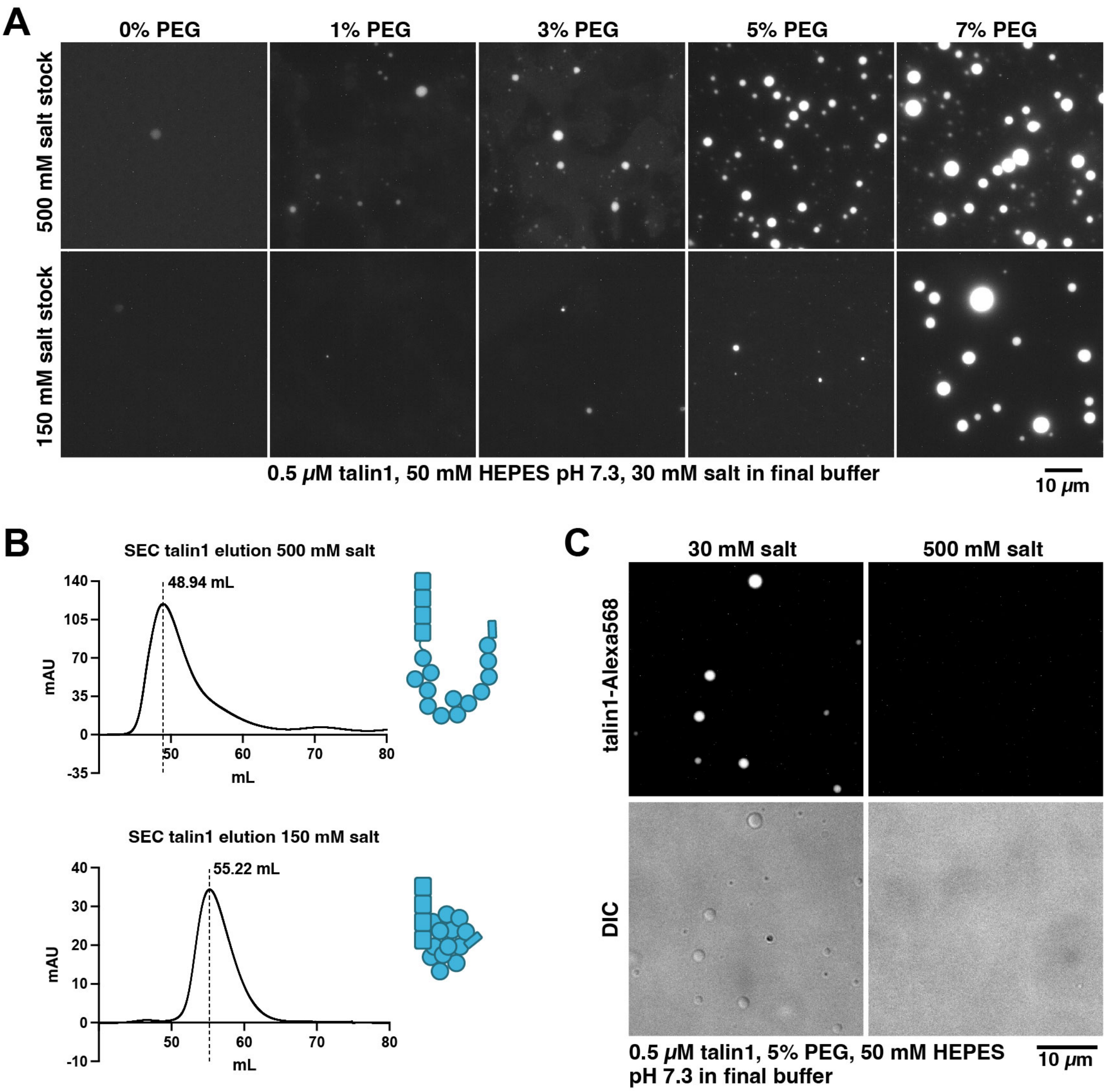


**S10. Talin1 phase separation can be induced by drastic reduction of salt in buffer.** Previous study has shown with cryoEM (Dedden et al 2019) that 80% of talin1 exhibit open conformation in buffer with 500 mM salt (NaCl or KCl), while only 20% are open in buffer with 150 mM salt. (A) Upon dilution of talin1 that was incubated at 500 mM salt (top panel) to low salt buffer (30 mM salt), phase separation occurs, relative to the amount of crowding agent (PEG) present. A similar trend is observed with talin incubated at 150 mM salt (bottom panel), although phase separation requires a much higher concentration of crowding agent, consistent with a smaller percentage of talin1 in an open conformation. (B) Talin1 elution from size-exclusion column with buffer containing different salt concentrations. With 500 mM salt, talin1 elutes earlier than with 150 mM salt, consistent with a more open, less globular conformation. (C) Talin1 condensates formed in the presence of crowding agent can be dissolved with high salt that disrupts protein-protein interaction.

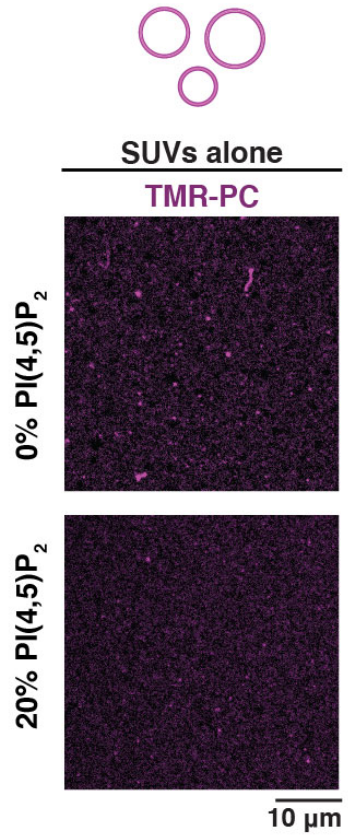


**S11. Control images for vesicles alone.** Under the same conditions as Fig. 3A without Tn2, SUVs are too small to be discernable. No droplets are observed for either lipid composition without additional proteins. Intensity has been increased relative to Fig. 3A in order to see lipid signal. Liposomes contained 74.75-X% DOPC, 15% DOPE, 10% DOPS, X% PI(4,5)$P_2$, 0.25% TopFluor-TMR-PI(4,5)$P_2$. Scale bar is 10 $\mu$m. All lipid-based experiments were carried out in the following buffer (10 mM HEPES pH 7.5, 100 mM NaCl).

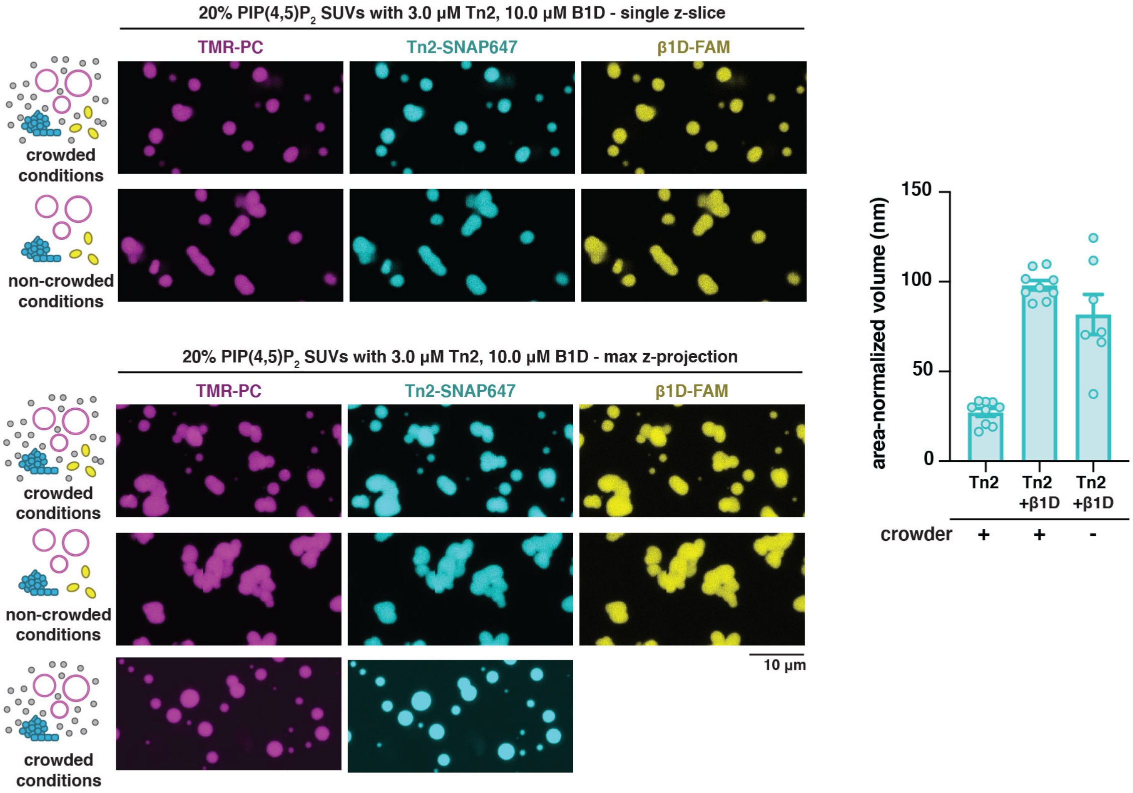


**S12. Tn2-β1D-SUV droplets form even in the absence of crowder. (A)** Single slice images of Tn2-β1D-SUV droplets with or without methyl cellulose. **(B)** Max z-projections from z-stacks of Tn2-β1D-SUV droplets with or without methyl cellulose. Droplets formed in the presence of all three components are less spherical, presumably because they do not reform a spherical shape after fusing. Instead, the droplets take on a pearls-on-a-string appearance, suggesting a gel-like condensate. The amount of phase separated material is much higher in the presence of both SUVs and β1D, regardless of whether crowder is included. Data represents pooled data from 3 independent samples for each condition. Error bars represent SEM. Liposomes contained 54.75% DOPC, 15% DOPE, 10% DOPS, 20% $PI(4,5)P_2$, 0.25% TopFluor-TMR-$PI(4,5)P_2$. All lipid-based experiments were carried out in the following buffer (10 mM HEPES pH 7.5, 100 mM NaCl), either with 0.25% methyl cellulose or without crowding reagent. Scale bar is 10 µm.

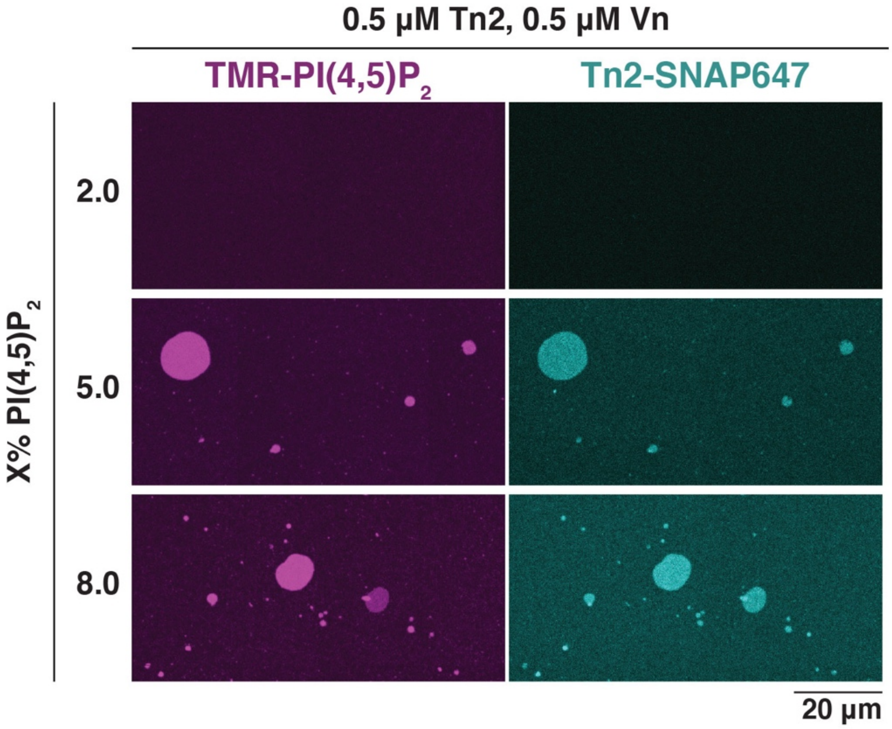


**S13. PI(4,5)$P_2$ drives formation of Tn2 clusters on SLBs.** Tn2 forms clusters on 5% and 8% PI(4,5)$P_2$ SLBs, but not on those with only 2% PI(4,5)$P_2$, suggesting there is a threshold for cluster formation dependent on membrane composition. As Tn2 membrane binding is dependent on PI(4,5)$P_2$ levels, this suggests that a minimum density of Tn2 molecules on the membrane surface is required for cluster formation. Proteins were mixed and added to the SLB immediately after bilayer formation. Samples were incubated for 1hour before imaging. Representative images are from experiments done in triplicate. SLBs were made from liposomes containing 74.75-X% DOPC, 15% DOPE, 10% DOPS, X% PI(4,5)$P_2$, 0.25% TopFluor® TMR PI(4,5)$P_2$. All lipid-based experiments were carried out in the following buffer (10 mM HEPES pH 7.5, 100 mM NaCl). Scale bar is 20 µm.

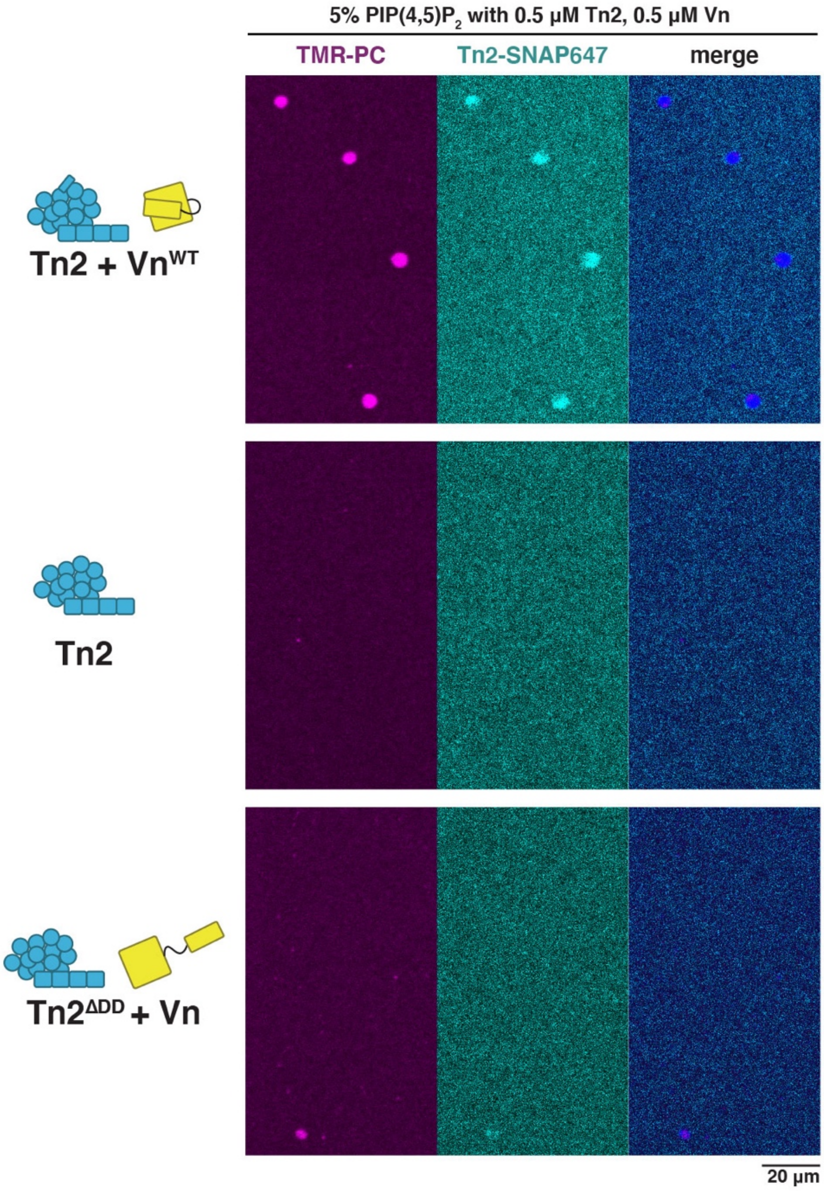


**S14 Tn2-PI(4,5)$P_2$ clusters on SLBs require Vn and the Tn2 dimerization domain.** Despite Tn2 binding to the membrane, no clusters are observed in the absence of Vn or $Tn2^{\Delta DD}$ with Vn. Proteins were mixed and added to the SLB immediately after bilayer formation. Samples were incubated for 1hour before imaging. SLBs were made from liposomes containing 69.75% DOPC, 15% DOPE, 10% DOPS, 5% PI(4,5)$P_2$, 0.25% TopFluor® TMR PI(4,5)$P_2$. Scale bar is 20 μm.

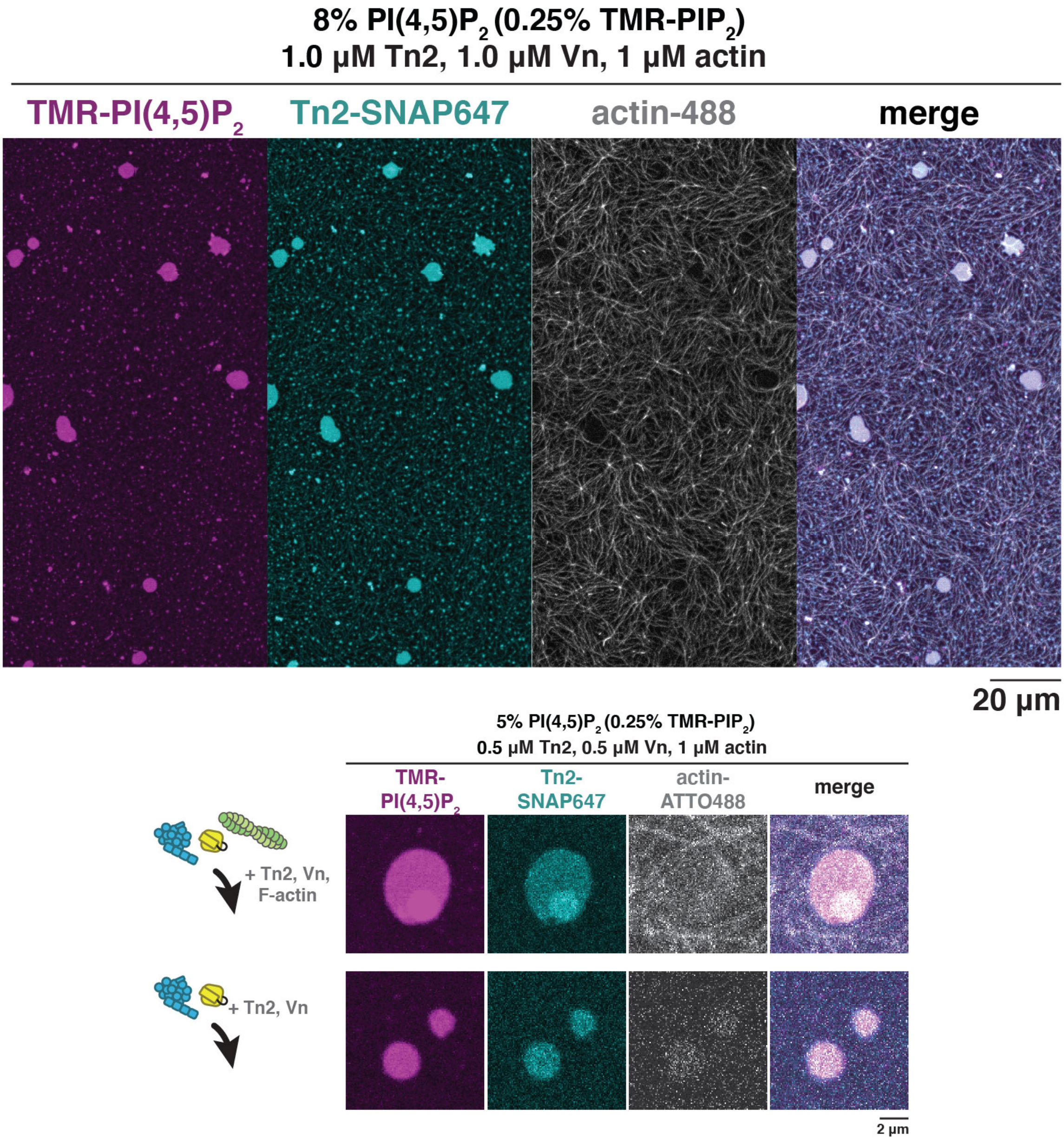


**S15. Actin is not recruited to Tn2-PI(4,5)$P_2$ clusters.** F-actin is recruited to the membrane surface in the presence of Tn2 and Vn, but does not become enriched within Tn2-PI(4,5)$P_2$ clusters. Actin was polymerized at RT for 30 minutes, then mixed with Tn,Vn at the indicated concentrations and incubated for 1 hr before imaging. A field-of-view image indicates that actin filaments localize around Tn2-PI(4,5)$P_2$ on the membrane, while the zoomed in view confirms that actin is not enriched within the clusters. Scale bars are indicated in the figure. SLBs were made from liposomes containing 66.75% DOPC, 15% DOPE, 10% DOPS, 8% PI(4,5)$P_2$, 0.25% TopFluor® TMR PI(4,5)$P_2$.

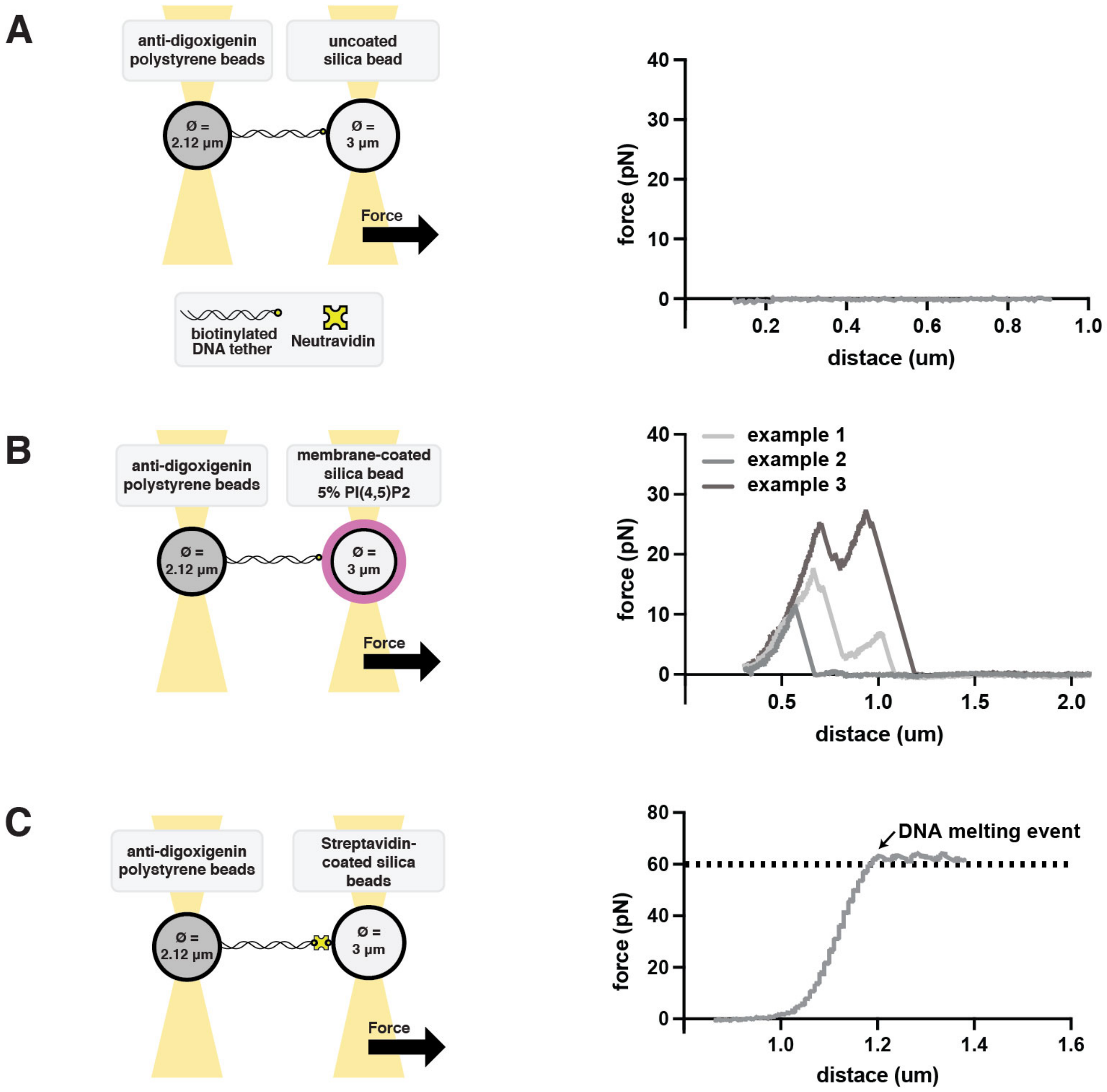


**S16. Tethering controls without talin.** (A) Tethers were not detected between anti-DIG beads with biotinylated DNA strands and uncoated silicon beads. (B) Some tethers were detected between anti-DIG beads with biotinylated DNA strands and silica beads coated with 5%PI(4,5)$P_2$ lipid membranes, but at a lower frequency than in the presence of talin. Additionally, the distance at which rupture occurred distinguished these tethers from true talin-membrane interactions, further described in SF17. Three example curves are shown, rupturing at various forces, but all below 1um distance. (C) A streptavidin-coated silica bead is used to demonstrate a stereotypical force curve for a single DNA strand between two beads, and the melting event that occurs at a force of 60 pN. (D) A worm-like chain model of a DNA force curve. (E) A worm-like chain model of a DNA-protein force curve, without unfolding events.

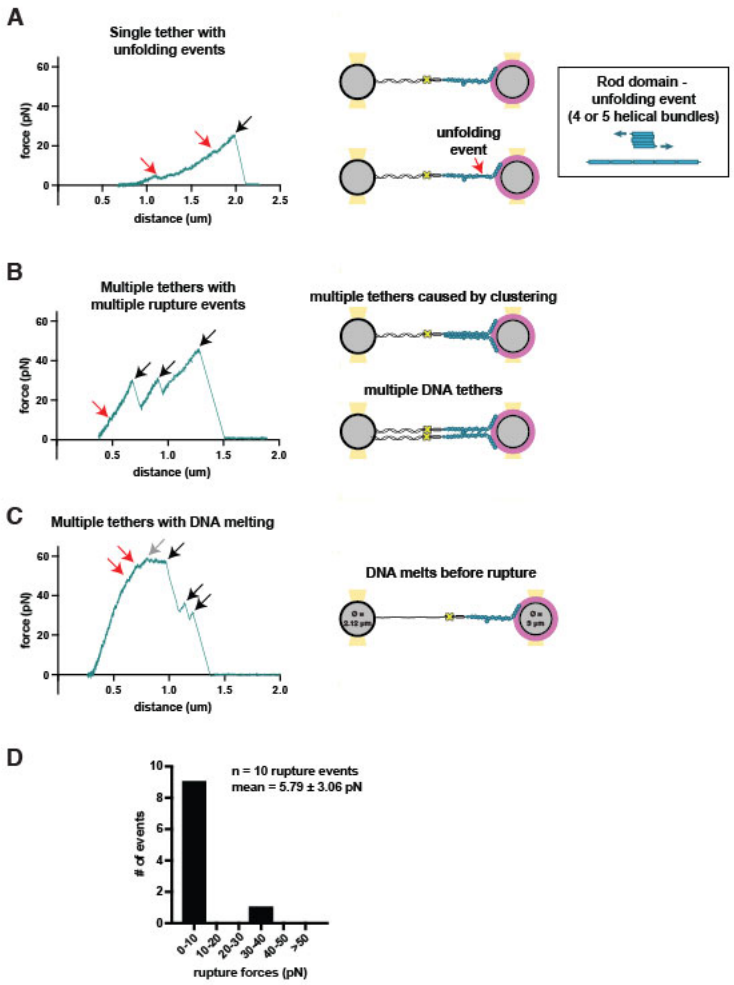


**S17. Examples of Tn2 tethers with 5% $PIP_2$.** Individual examples of force curves obtained with Tn2 bound to 5% PI(4,5)$P_2$-containing membrane-coated beads. (A) A single DNA-Tn2 tether, with multiple unfolding events and a rupture event at a distance of 2 µm. The inset figure shows a model of an individual helical bundle within the talin rod domain. These helical bundles are made up of either 4 or 5 helices. When placed under tension, as shown here, the helical bundle unfolds, taking on an extended, linear conformation. (B) An example of multiple Tn2-tethers pulled simultaneously, and rupturing sequentially. The steep initial increase in force is indicative of multiple tethers, which could be due to multiple Tn2-DNA tethers, or due to interactions between the membrane-bound talin molecules themselves. (C) An example of multiple DNA-Tn2 tethers which withstand more than 60 pN of force, at which point the DNA strand melts. This is an example of a connection too strong to be included in analysis.

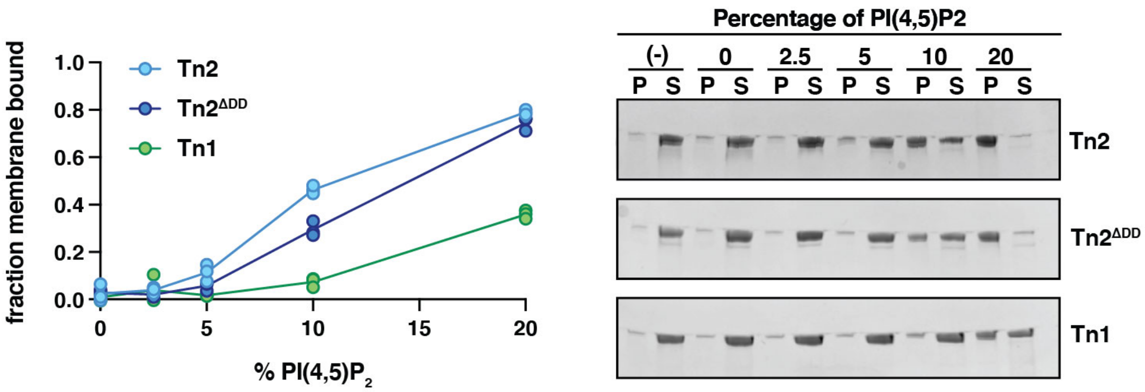


**S18. Tn2, Tn1, and Tn2$^{\Delta DD}$ membrane binding depends on PI(4,5)$P_2$ levels**. Tn2 and Tn2$^{\Delta DD}$ bind at similar levels, while Tn1 cosediments at lower levels even with the highest amount of PI(4,5)$P_2$. This is consistent with published reports that Tn1 has a lower affinity for membrane binding. The graph shows the fraction of total protein present in the pellet after centrifugation with vesicles with the indicated percentage of PI(4,5)$P_2$. Each condition was carried out in triplicate in buffer of 20 mM HEPES, pH 7.5 and 100 mM NaCl. Representative SDS-PAGE gels are shown on the right, with pellet (P) and supernatant (S) samples for each protein at each lipid composition, including a no lipid control sample (-). Liposomes contained 75-X% DOPC, 15% DOPE, 10% DOPS, X% PI(4,5)$P_2$. Each condition represents data from three independent incubations.
