## Supplementary material for "Membrane-induced 2D phase separation of focal adhesion proteins": Materials and Methods

### Plasmids

Plasmids for Talin2, Vn, $Vn^{3Q}$ and $Vn^{DR}$ were previously made for Kelley et al. (*1*). We refer to these here as pET21_hTalin2 L435G -3C-SNAP-His, pET21_hVinculin-3C-SNAP-His, pET21_hVinculin K944Q R945Q K1061Q-SNAP-His and pET21_hVinculin K944Q R945Q K1061Q-SNAP-His. All Talin2 plasmids contain the L435G mutation to reduce calpain cleavage.

To generate plasmids for expressing Talin1 as his-SNAP-tagged fusions in E. coli, ORFs were PCR amplified from pET101-hTalin1-His (*2*) and assembled with the PCR amplified backbone from pET21_hTalin2 L435G -3C-SNAP-His (*1*) using seamless cloning (ThermoFisher Scientific/Invitrogen™ GeneArt™ Seamless Cloning and Assembly Enzyme Mix), resulting in pET21_hTalin1-3C-SNAP-His.

Site-directed mutagenesis was performed on pET21_hTalin2 L435G-3C-SNAP-His (*1*) to generate the S339L and E1772A mutants and assembled with homologous recombination or Seamless Cloning (GeneArt™) respectively, resulting in pET21_hTalin2 L435G E1772A-3C-SNAP-His and pET21_hTalin2 L435G S339L-3C-SNAP-His.

The plasmid for the ΔDD Tn2 truncation was amplified from pET21_hTalin2 L435G-3C-SNAP-His (*1*) resulting in pET21_hTalin2 L435G 1-2493 -3C-SNAP-His.

For $Vn^{DR}$, site directed mutagenesis was performed on pET21_hVinculin K944Q R945Q K1061Q-SNAP-His (*1*) to introduce the additional N773A and E775A mutations and then assembled using Gibson Assembly (*3*), resulting in pET21_hVinculin N773A E775A K944Q R945Q K1061Q-SNAP-His. Site directed mutagenesis then also was performed on the resulting plasmid to introduce the A50I mutation and assembled with homologous recombination, resulting in pET21_hVinculin A50I N773A E775A K944Q R945Q K1061Q-SNAP-His.

The plasmid for the $Vn^{D1}$ fragment was amplified from pET21_hVinculin-3C-SNAP-His (*1*) and assembled by blunt-end cloning, resulting in pET21_hVinculin 1-258 -3C-SNAP-His.

### Protein expression and purification

Constructs were expressed in *E. coli* BL21 (DE3) gold using ZY auto-induction medium. Talin proteins were all purified using the same protocol, based on that described in a previous report (*2*). Cells were lysed by sonication in 50 mM Tris-HCl pH 7.8, 500 mM NaCl, 5 mM imidazole, 3 mM β-mercaptoethanol, 1 mM EDTA, and Roche cOmplete protease inhibitor tablets (Roche, Basel, Switzerland), followed by purification using nickel-affinity chromatography (cOmplete His-Tag purification column, Roche), and cation exchange (HiTrap SP FF, GE Healthcare, Chicago, Illinois). Next, the his-tag was either removed using overnight incubation with 3C protease, or labeled using overnight incubation with SNAP-AlexaFluor647 (New England Biolabs, Ipswich, Massachusetts). Finally, protein was further purified by size-exclusion chromatography using either a Superdex 200 16/600 column (GE Healthcare) or Superose 6 10/300 column (GE Healthcare) in 50 mM HEPES pH 7.8, 150 mM KCl, 3 mM β-mercaptoethanol, 1 mM EDTA, and 10% glycerol, followed by flash freezing for storage at −80°C.

For vinculin proteins, cells were lysed by sonication in 50 mM Tris-HCl pH 7.8, 500 mM NaCl, 5 mM imidazole, 3 mM β-mercaptoethanol, 1 mM EDTA, and Roche cOmplete protease inhibitor tablets. Following lysis, TritonX-100 was added for a final amount of 1% by volume. Full-length vinculin cell lysates were incubated on Roche cOmplete His-Tag resin for 2 hr at

4°C, then washed with 50 mM Tris-HCl pH 7.8, 500 mM NaCl, 10 mM imidazole, 3 mM β-mercaptoethanol, 1 mM EDTA. After washing, proteins were incubated overnight with either 3C protease to remove the SNAP-his tag, or labeled with SNAP-AlexaFluor488 or SNAP-Surface594 (New England Biolabs). Following removal or elution from beads, vinculin proteins were then further purified by size-exclusion chromatography using Superdex 200 16/600 column (GE Healthcare) or Superose 6 10/300 column (GE Healthcare) in 50 mM HEPES pH 7.8, 150 mM KCl, 3 mM β-mercaptoethanol, 1 mM EDTA. Vinculin fragments were purified using nickel-affinity chromatography, immediately eluted from the column with 1M imidazole, cleaved overnight with 3C, and separated from the cleaved SNAP-his tag by reverse nickel-affinity chromatography. This was followed by size-exclusion chromatography using a Superdex 75 10/300 in 50 mM HEPES pH 7.8, 150 mM KCl, 3 mM β-mercaptoethanol, 1 mM EDTA. Proteins were flash frozen and stored at −80°C.

Actin proteins were purchased in lyophilized form from HYPERMOL.

#### β1D Peptide

The fluorescently labeled β1D peptide was produced by the Bioorganic Chemistry & Biophysics Core Facility at the Max-Planck-Institute of Biochemistry by solid phase synthesis using Fmoc/*t*Bu chemistry and microwave heating on a Liberty Blue peptide synthesizer (CEM Corporation, Mathews, NC, U.S.A.).

#### Confocal microscopy of biomolecular condensates

For most experiments were performed in flow chambers. For this, glass coverslips were washed extensively with milliQ water and dried with nitrogen gas, then coated using a solution of 80% ethanol pH 2, 2 mg/mL methoxy-poly (ethylene glycol)-silane and dried for several hours at 75°C. Immediately before using, coverslips were washed extensively with milliQ water, dried under a stream of nitrogen, and attached to adherent flow chambers (Ibidi, Martinsried, Germany).

For the experiments in Figure S4 we used microtiter plates (Greiner Bio-One, 384-well glass bottom SensoPlate™), which we passivated beforehand with 50 µl of 5 mg/ml β-casein (Sigma Aldrich) for 20 min.

Imaging was performed with a Zeiss LSM 780/CC3 confocal microscope equipped with a C-Apochromat, 63x/1.4 W objective. PMT detectors (integration mode) were used to detect fluorescence emission (excitation at 488 nm for ATTO488, 594 nm for SNAP594 and 633 nm for SNAP647) and record confocal images. All experiments were conducted at room temperature.

#### Quantification

The amount of phase separated material in confocal microscope data was quantified with a custom written code. These results are shown in Figures 1C, 2D and 3B and supplementary Figures S4, S6, S7, S8 and S12. Our Matlab code first binarizes all z-slices of a confocal z-stack and determines the area of biomolecular condensates visible in each z-slice. The volume is then extrapolated by just adding up the areas of all z-slices, which are spaced 1 µm apart. We then divide the resulting volume by the total area of the field of view, to get an area-normalized volume, i.e. the average height of a droplet, equivalent to how most types of precipitation is quantified.

### Analytical size-exclusion chromatography assays

Proteins used were first buffer exchanged or diluted to match the conditions tested (i.e. 75 mM or 500 mM KCl). Proteins were incubated together on ice for 15 min, and prespun in a tabletop microcentrifuge at 15,000 x g. 75% of the sample volume was removed, avoiding any potential pellet, and applied to a Superose 6 Increase 3.2/300 column with 20 mM HEPES pH 7.8, 1 mM EDTA, 3 mM β-mercaptoethanol, and either 75 mM or 500 mM KCl.

### Supported lipid bilayer assays

The SLBs were prepared in the same fashion as we described previously (*1*) and is based on a protocol by Braunger et al. (*4*). Briefly, lipids were mixed in a glass vial, and dried under a continuous stream of $N_2$, then dried overnight in a vacuum chamber at room temperature. The lipid film is then gently rehydrated in citric acid buffer pH 4.8, and incubated at room temperature for 20 min before vortexing briefly. To produce small unilamellar vesicles (SUVs), the solution is then sonicated for 30 min (30 s on/30 s off intervals).

Coverslips were cleaned using piranha solution for at least 15 min. Immediately before using, coverslips were rinsed thoroughly in deionized water and dried with $N_2$, then attached to Ibidi adherent flow chambers. To form SLBs, 60 µL of 0.2 mg/mL SUVs were added to individual flow chambers and incubated for 3 min, followed by 2 × 80 µl imaging buffer buffer (1x) to remove excess vesicles.

SLBs were then tested for bilayer integrity and fluidity using fluorescence recovery after photobleaching (FRAP). To the fluid bilayers, proteins were added in liposome buffer supplemented with GODCAT.

For experiments on SLBs that include actin, we preincubate the SLBs with 0.5 mM Tn2 for 15 minutes. To the fluid bilayers, proteins and actin (5% ATTO488-actin) were added in imaging buffer supplemented with GODCAT.

Imaging was performed with an LSM 780/CC3 confocal microscope (Carl Zeiss, Germany) equipped with a C-Apochromat, 40×/1.2 W objective. PMT detectors (integration mode) were used to detect fluorescence emission (excitation at 488 nm for ATTO488, 594 nm for SNAP594 and 633 nm for SNAP647) and record confocal images.

### Lipid co-sedimentation assays

Lipid co-sedimentation assay were conducted as previously described (*5, 6*). Briefly, liposomes were swelled from dried lipid in 20 mM HEPES, pH 7.5 and 100 mM NaCl. FA proteins were mixed with 1 mg/mL liposomes and incubated at room temperature for 30 min, then spun at 18000 x g in a tabletop microcentrifuge at 4°C. Equal volumes of pellet and supernatant were analyzed by gradient SDS-PAGE, and quantified using Fiji. For quantification, the percent of protein in the pellet of protein-alone control samples was subtracted from all experimental samples.

### Optical tweezers assay to study talin-membrane interactions

To extract forces acting between talin and the $PIP_2$-doped lipid bilayer, we designed an assay using two optically trapped beads, one of which was functionalized with the $PIP_2$-doped lipid bilayer and talin, and the other was functionalized with short DNA handles binding to biotinylated talin structures via neutravidin (see Figure 4A). A similar assay to probe protein-membrane interactions was previously used by Ma et al. (*7*).

### Preparation of functionalized beads

Membrane-coated beads were prepared by spreading small unilamellar vesicles of defined lipid composition on silica beads as described earlier (*8*). Briefly, dried lipid films are prepared and equilibrated at RT. 500 µl of buffer are added to 0.2 mg lipids, and incubated for 30 min at RT to let the films swell, vortexed until all lipid material was in suspension and sonicated for 8 x 1 min in ice with 1 minutes breaks until the suspension was opaque. We then centrifuged the suspension at 4°C and 15000g for 5 min, to separate multilamellar material from unilamellar bilayers. We used the supernatant/unilamellar fraction to coat the silica beads. 70 µL of silica beads (Spherotech SIP-30-10 Sphero Silica Particles, 3 µm, 5% w/v) were cleaned by 3 centrifuging/washing cycles with 50 µL of buffer each (final volume approx. 70 µL). Cleaned beads (18 µl) were incubated with 220 µL vesicle suspension, vortexed for 45 min to shear additional vesicle material off the membrane-coated beads and spinned them down at 500g for 1 h. Supernatant was removed and the pellet was resuspended with 500 µL measurement buffer and washed 2 times before using the membrane-coated bead suspension for the measurement. Coated beads were functionalized with talin by incubating 6 µL of bead solution with 2 µL of 0.5 µM Talin solution for 1 min at RT. After incubation, the bead suspension was diluted to 1 mL with measurement buffer and used for the experiment.

Beads with DNA handles were prepared by incubating 1 µL of anti-digoxigenin functionalized polystyrene beads (Spherotech Anti-dig-coated polystyrene beads, 2.12 um, 0.1 w/v%) with 1 µL 3kb DNA handles (3xdig/3kb DNA/3xbiotin handle, approx. 20 ng/mL).The functionalized beads were then incubated with 3 µl of 1 mg/mL neutravidin solution for 1 min. After incubation, the bead suspension was diluted to 1 mL with measurement buffer and used for the experiment.

### Setting up the optical tweezers assay

To setup the assay, we used a LUMICKS flow cell and microfluidic system in a confocal C-trap setup (LUMICKS) that allows for recording force and fluorescence data in parallel.

To probe protein-membrane interaction forces, we used the following workflow: first, we trapped a membrane-coated bead in one trap and a DNA handle-functionalized bead in the second trap. We used the confocal scanning modality of the instrument to confirm that the fluorescently labelled lipid bilayer is homogeneously spread on the silica beads.

We established the binding between DNA handle to talin on the membrane-coated beads by approaching the two beads. Subsequently, we recorded force-extension curves (FEC) by moving the bead in the second trap away from the bead in the first trap and read out interaction forces acting between talin and membrane. Since rupture forces typically depend on the speed at which the optical trap moves, we ensured a constant speed of 0.25 um/s throughout the experiments.

Before studying the full protein construct, we ran a sequence of control measurements (Figure S16). We tested

i) the 3kb DNA handle by directly binding it to one anti-digoxigenin and one streptavidin-coated bead and recording typical DNA melting curves. We optimized the DNA to having a single DNA handle between the beads.

ii) the interaction of the DNA handle with a 0% PIP2 lipid bilayer, as well as the

iii) the interaction of the DNA handle streptavidin construct with the 0% and 5% PIP2-doped membrane without the protein.

**Analysis of optical tweezers data**

Before analyzing FECs, we first subtracted the background force: we recorded an FEC without tether and subtracted that from the final data. High resolution distance data (all force and trap data were recorded at 78 kHz) were obtained by looking at the trap-trap distance and subtracting the bead displacement. For plotting purposes, we downsampled the FEC by a factor of 500. An example code showing how we obtained the high-resolution data and subtracted the background can be found here: https://harbor.lumicks.com/single-script/b7b98127-1b09-4505-9967-ff1c0b2aaf92.

We were interested in the rupture force of a single DNA tether and one (or more) talin complexes bound to the membrane. Therefore, the protocol for extracting rupture forces from FECs was as follows: 1) If the tether had multiple force jumps, we would only extract the value of the last jump, when the force jumps to zero, indicating that the protein handle is fully detached from the membrane. 2) We monitored not only the force at which a rupture occurs, but also the distance. The distance allowed us to compare various constructs. 3) We ignored rupture force values above 60 pN. A single strand of dsDNA melts at 60 pN, so rupture forces above this value would indicate the presence of multiple DNA handles bound to the functionalized membrane.
